# Two-pass alignment using machine-learning-filtered splice junctions increases the accuracy of intron detection in long-read RNA sequencing

**DOI:** 10.1101/2020.05.27.118679

**Authors:** Matthew T. Parker, Katarzyna Knop, Geoffrey J. Barton, Gordon G. Simpson

**Affiliations:** School of Life Sciences, University of Dundee, Dow Street, Dundee, DD1 5EH, UK; James Hutton Institute, Invergowrie, DD2 5DA, UK

**Keywords:** splicing, long read sequencing, spliced alignment, RNA-seq, gene expression, transcriptome assembly, machine learning, nanopore

## Abstract

Transcription of eukaryotic genomes involves complex alternative processing of RNAs. Sequencing of full-length RNAs using long reads reveals the true complexity of processing. However, the relatively high error rates of long-read sequencing technologies can reduce the accuracy of intron identification. Here we apply alignment metrics and machine-learning-derived sequence information to filter spurious splice junctions from long read alignments and use the remaining junctions to guide realignment in a two-pass approach. This method, available in the software package 2passtools (https://github.com/bartongroup/2passtools), improves the accuracy of spliced alignment and transcriptome assembly for species both with and without existing high-quality annotations.

## Background

Understanding eukaryotic genomes requires knowing not only the DNA sequence but also which RNAs are transcribed from it. Eukaryotic transcription by DNA-dependent RNA polymerase II is associated with multiple alternative RNA processing events that diversify the coding and regulatory potential of the genome. Alternative processing choices include distinct transcription start sites, the alternative splicing of different intron and exon combinations, alternative sites of cleavage and polyadenylation, and base modifications such as methylation of adenosine. Patterns of alternative processing can be extensive. For example, more than 90% of human protein-coding genes have at least two splice isoforms(1). Changes in RNA processing can reflect the reprogramming of gene expression patterns during development or in response to stress or result from genetic mutation or disease. Consequently, the identification and quantification of different RNA processing events is crucial to understand not only what genomes encode but also the biology of whole organisms(2).

The sequencing of RNAs (RNAseq) can reveal gene expression patterns in specific cells, tissues or whole organism. The success of this approach depends upon sequencing methodology and the computational analyses used in interpreting the sequence data. High-throughput sequencing of RNA rarely involves direct RNA sequencing (DRS): instead, copies of complementary DNA (cDNA) produced by reverse transcription of RNA molecules are sequenced(2). However, template strand switching by reverse transcriptase (RT) during the copying process can produce spurious splicing patterns and antisense RNA signals(3, 4). Three current technologies use RT-based RNA sequencing library preparation: Illumina, Pacific Biosciences (PacBio) and Oxford Nanopore Technologies (ONT). Illumina RNAseq can generate hundreds of millions of highly accurate short sequencing reads, each representing a 50–250 nt fragment of full-length RNA(2). Methods exist for quantifying known alternative splicing events from short reads(5). However, when the transcript models are unknown, for example in a non-model organism or a mutant or disease with altered RNA processing, new transcript models must be generated, either *de novo* or with the aid of the reference genome. Because Illumina reads are short, they are unlikely to overlap multiple splice junctions, meaning that phasing of splicing events is difficult and requires complex computational reconstruction(6–8). PacBio and ONT can sequence full-length cDNA copies without fragmentation, thus allowing whole transcript isoforms to be identified unambiguously(2). Most recently, ONT introduced a direct sequencing method for RNA(9–11). Using this approach, it is now possible to capture information on the splicing, 5′ and 3′ ends, poly(A) tail length, and RNA modifications of full-length RNA molecules in a single experiment, without RT-associated artefacts(11).

The development of technologies for sequencing full-length RNA molecules makes the identification of authentic processing events possible in principle, but software tools are also needed to interpret the RNA processing complexity. PacBio and ONT sequencing reads have a higher error rate than Illumina(10–14). Consequently, alignment accuracy for long sequence reads at splice junctions is often compromised(9–11). This is a problem for genome-guided transcriptome annotation because the incorrect identification of splice junctions leads to mis-annotated open reading frames and incorrectly truncated protein predictions. In addition, if alignment errors are systematic (i.e. occur for transcripts with specific characteristics), then quantification of transcripts will be compromised. Even with completely error-free reads, alignment at splice junctions is often confounded by multiple equally plausible alternatives(15). Accordingly, computational methods for improving the splice-aware alignment of long reads are required.

Software tools for long and short RNAseq data analysis incorporate several approaches to address the challenges presented by pre-mRNA splicing. Biologically relevant information can aid the alignment of transcriptomic sequences to the genome. For example, the vast majority of eukaryotic splicing events occur at introns bordered by GU and AG motifs. Making RNAseq read aligners aware of these sequence features (as is the case for the commonly used spliced aligners STAR(16), HISAT2(17) and minimap2(18)) can significantly improve the alignment of reads at splice junctions. In addition, where genome and transcriptome annotations exist, many alignment tools allow users to provide sets of correct splice junctions to guide alignment(16–19). Introns containing these guide splice junctions are penalised less than novel introns, resulting in fewer alignment errors. For long reads, software tools such as FLAIR(10) use post-alignment correction to improve splice junction detection and quantification. Post-alignment correction tools take long-read alignments and guide splice junctions from either a reference annotation or a set of accurate short RNAseq reads(10). Introns from long-read alignments which are not supported by the guide splice junction set are “corrected” to the nearest supported junction within a user-defined range. It is unclear whether such post-alignment corrections confer any benefit over providing guide splice junctions during alignment. Small errors in spliced alignment can also be corrected during reference-guided transcriptome assembly. Tools such as StringTie2(6) and pinfish (Oxford Nanopore Technologies) identify clusters of similarly aligned reads and correct them to the median junction positions, before outputting annotations.

Two-pass alignment has also been used to improve splice junction detection and quantification(16, 19, 20). In a two-pass alignment approach, splice junctions detected in a first round of alignment are scored less negatively in a second round, thereby allowing information sharing between alignments. This approach has been useful for short-read data, where RNA fragmentation may occur close to splice junctions during sequencing library preparation. The two-pass approach enables these short junction overhangs to be aligned to splice junctions detected in other alignments(20). Splice junctions detected in a first pass may also be filtered to remove false positives before second-pass alignment. Existing tools for splice junction filtering, such as finesplice and portcullis(21, 22), use machine learning with training on a range of junction metrics. A model is trained from high-confidence positive and negative examples from training data and then applied to classify the remaining splice junctions at the decision boundary. Splice junctions are then filtered to remove junctions predicted to be spurious. Subsequent second-pass alignment guided by these filtered junctions can then improve the accuracy of alignment(22).

In this study, we develop a method for filtered two-pass alignment of the relatively high-error long reads generated by techniques such as nanopore DRS. The resulting software, which we have named 2passtools, uses a rule-based approach to identify probable genuine and spurious splice junctions from first-pass read alignments. These can then be used to train a logistic regression (LR) model to identify the biological sequence signatures of genuine splice junctions. We found that integrating the alignment and sequence information extracted in this manner produced the largest improvement in splice junction alignment and subsequent genome-guided annotation. As a result, we can improve the utility of long-read sequencing technologies in revealing the complexity of RNA processing and annotating newly sequenced organisms.

## Results and Discussion

### Reference-splice-junction-aware alignment is more accurate than post-alignment junction correction

For sequencing experiments designed to interpret RNA from model organisms, a set of reference splice junctions will already be available (e.g. from Ensembl). We therefore asked how providing these reference splice junctions to minimap2 to guide alignment performed compared with post-alignment correction of junctions with FLAIR(10). For this analysis, we used four nanopore DRS datasets generated from Arabidopsis seedlings(11) and four datasets generated from human cell lines(10). Several types of probable alignment error were identifiable in these data, including failure to align terminal exons and short internal exons, spurious terminal exons, and large insertions to the reference genome (Fig. 1). Because these datasets are likely to contain novel splice junctions which do not appear in reference annotations, we simulated full-length reads (i.e. with no 3′ bias(11)) using the Arabidopsis and human reference transcriptomes, AtRTD2(23) and GRCh38(24), respectively. Simulated reads were then mapped to the corresponding reference genome using minimap2(18), either with or without guidance from reference splice junctions. Alignments of simulated reads were found to have similar error profiles to genuine nanopore DRS read alignments (Fig. S1). Reads mapped without reference splice junctions were then corrected using FLAIR with reference splice junctions.

**Fig. 1.**
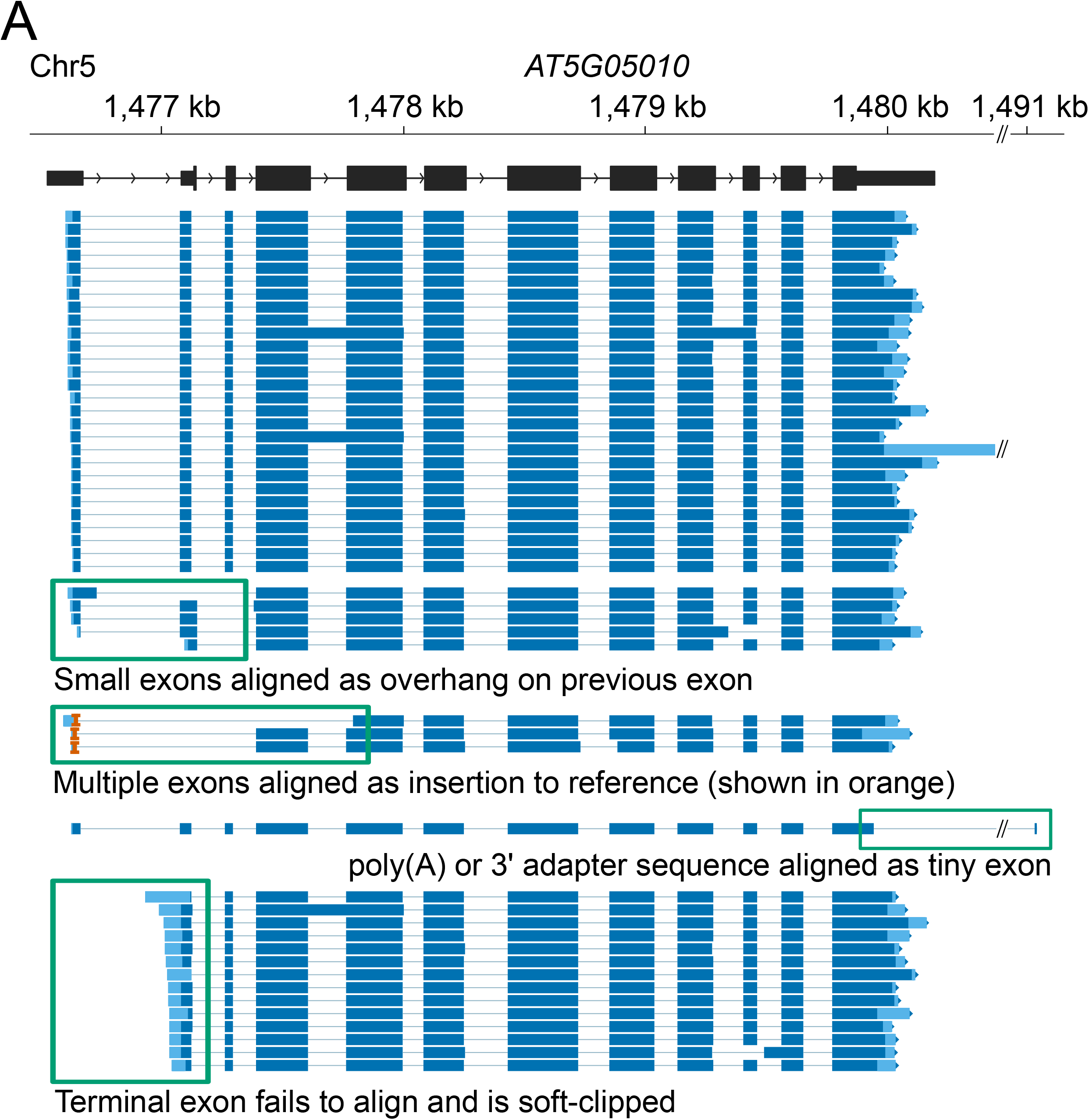
Assessment of alignment errors in nanopore DRS datasets. Nanopore DRS read alignments at Arabidopsis *AT5G05010* locus with different types of alignment error presented. Read alignments are shown in dark blue, with soft-clipped (unaligned) ends shown in light blue. Mismatches and indels of <30 nt are not shown. Insertions to the reference of > 30 nt are shown as orange carets.

Although nanopore DRS has some systematic errors in base-calling (particularly at homopolymers), the majority of sequencing errors occur stochastically(25). In contrast, we found that alignment errors were often repeated at similar locations in the alignments of independent reads from equivalent mRNA transcripts (Fig. 1, Fig. 2A). A common alignment error at splice junctions is failure of a short exon to align correctly. Instead, fragments of the exon are aligned to the ends of flanking exons, resulting in a single incorrectly defined intron. A clear example of such an alignment error was detected at the short (42 nt) exon 6 of Arabidopsis *FLM* (*AT1G77080*; Fig. 2A). Minimap2 uses a modified form of the Smith– Waterman algorithm for performing local alignment(18, 26). This method scores alignments using bonuses for matches to the reference sequence and penalties for mismatches or the opening of insertions, including introns. Incorrect alignment of *FLM* exon 6 is likely to occur because the bonus for aligning a short exon with sequencing errors is not sufficient to overcome the penalty for opening the two flanking introns(18). Overall, we found that only 19.3% of simulated *FLM* reads aligned to the correct transcript isoform. Because the sequence distance between the alignment and the genuine reference splice junctions was so great, FLAIR was unable to perform post-alignment correction at *FLM* exon 6, resulting in the reporting of incorrect introns (Fig. 2A). In all, 40.3% of simulated *FLM* reads were aligned to the correct transcript isoform after FLAIR correction of splice junctions using the reference annotation. However, providing reference splice junctions to minimap2 during alignment resulted in the correct identification of *FLM* exons and introns in most cases: 92.1% of simulated *FLM* reads were aligned to the correct transcript isoform. We conclude that for loci with complex splicing patterns, reference-splice-junction-guided alignment performs better than post-alignment correction.

**Fig. 2.**
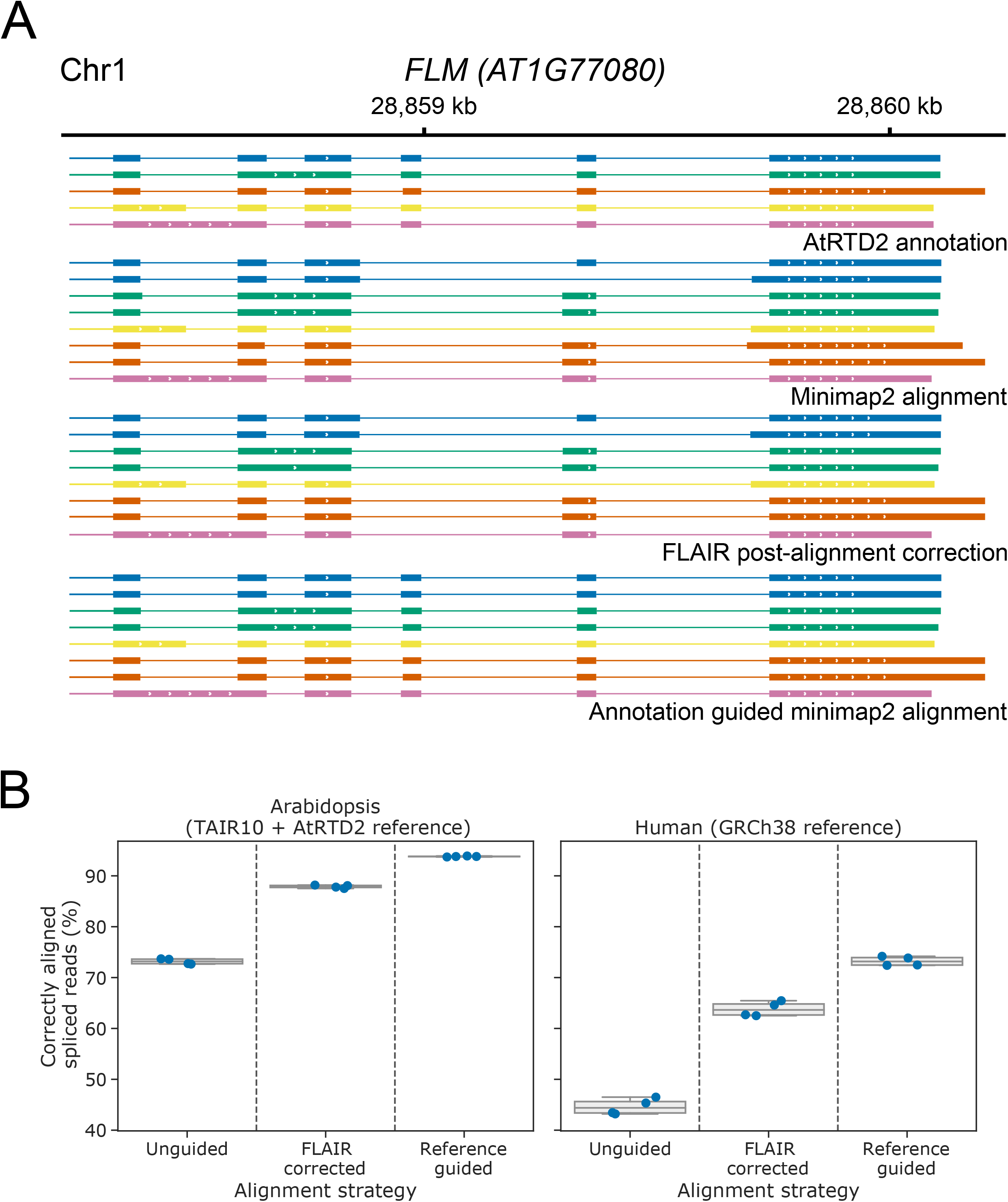
Improved spliced alignment of simulated reads using annotation-guided alignment. **A** Reference-guided alignment improves the identification of small exons in nanopore DRS reads. Gene track showing the alignment of a sample of simulated nanopore DRS reads at the Arabidopsis *FLM* gene. AtRTD2 reference annotation, from which reads were simulated, is shown on top, with unguided minimap2 alignments, FLAIR correction of unguided minimap2 alignments and reference-guided minimap2 alignments shown below. Only reads where exon 6 failed to align in the initial unguided alignment are shown. Each read alignment is coloured based on the reference transcript it was simulated from, and reads are shown in the same order within each alignment method group. Mismatches and indels are not shown. **B** Reference-guided alignment improves the identification of correct transcripts globally. Boxplots with overlaid strip-plots showing the percentage of alignments which map exactly to the splice junctions of the transcript from which they were simulated, for unguided minimap2 alignments, FLAIR correction of unguided minimap2 alignments using reference annotation, and reference annotation-guided minimap2 alignments. Reads simulated from intronless transcripts which map correctly without splicing were not included in percentage calculations. Reads were simulated from Arabidopsis (left) and human (right) nanopore DRS data aligned to the AtRTD2 and GRCh38 reference transcriptomes, respectively.

Without guidance from a reference annotation, we found that a median of 73.2% of Arabidopsis reads and 44.4% of human reads mapped correctly to the splice junctions of the transcript they were simulated from (Fig. 2B). The difference between the two organisms may be explained by biological differences between the two species (e.g. in intron size, number of exons per transcript, number of intronless transcripts). After post-alignment correction of splice junctions using FLAIR, the number of correctly identified transcripts detected was improved (median of 87.9% and 63.6% for Arabidopsis and human reads, respectively; Fig. 2B). This came at the cost of a small increase in alignment of reads to incorrect reference transcript splice junctions: from a median of 1.79% to 2.62% for Arabidopsis and from 3.86% to 5.45% for human (Fig. S2A). This misclassification may affect the relative quantification of transcripts for some genes, with implications for differential transcript usage analysis. Reference annotation-informed alignment with minimap2 performed better than FLAIR, with a median of 93.8% of Arabidopsis reads and 73.2% of human reads aligning correctly at the splice junctions of the transcript they were simulated from (Fig. 2B), albeit with misclassification rates of 2.61% and 5.49% respectively (Fig. S2A). We conclude that there is a clear benefit to providing reference splice junctions during alignment of long reads with relatively high sequence error rates, and that this is preferable to post-alignment correction.

### Alignment metrics enable identification of genuine splice junctions

In newly sequenced organisms, suitable reference annotations to guide alignment may not be available. We therefore asked how the spliced alignment of nanopore DRS reads might be improved in the absence of reference annotation. Naïve two-pass alignment has been successfully used to improve the spliced alignment of short reads(20). We applied this approach with our real and simulated nanopore DRS reads. Splice junctions identified by a first-pass alignment of reads were selected and used (without filtering) to inform a second-pass alignment. The method was compared with reference-guided alignment with minimap2, since we find this to be the gold-standard for aligning reads using information from a reference annotation. We found that using the naïve two-pass approach, the median percentage of simulated Arabidopsis DRS alignments which matched the splice junctions of the reference transcript they were simulated from could be increased slightly from 73.2% to 75.8% (Fig. S2B). The increase was similar for reads simulated from human DRS alignments: from 44.4% to 47.3% (Fig. S2B).

We next considered whether further improvements in two-pass alignment could be obtained by filtering out likely false-positive splice junctions from first-pass alignments. This would allow us to provide more refined guide junctions for second-pass alignment (Fig. 3A). A similar approach worked for short reads when splice junctions were filtered by using junction metrics to train a classifier in the portcullis software tool(22). By using the presence or absence of a splice junction in the reference annotation as a ground truth, we considered a range of novel or previously introduced junction metrics(21, 22), including junction alignment distance, supporting read count, intron motif and the presence/absence of nearby splice donor and acceptor sites with higher supporting read counts (Fig. S3A-D).

**Fig. 3.**
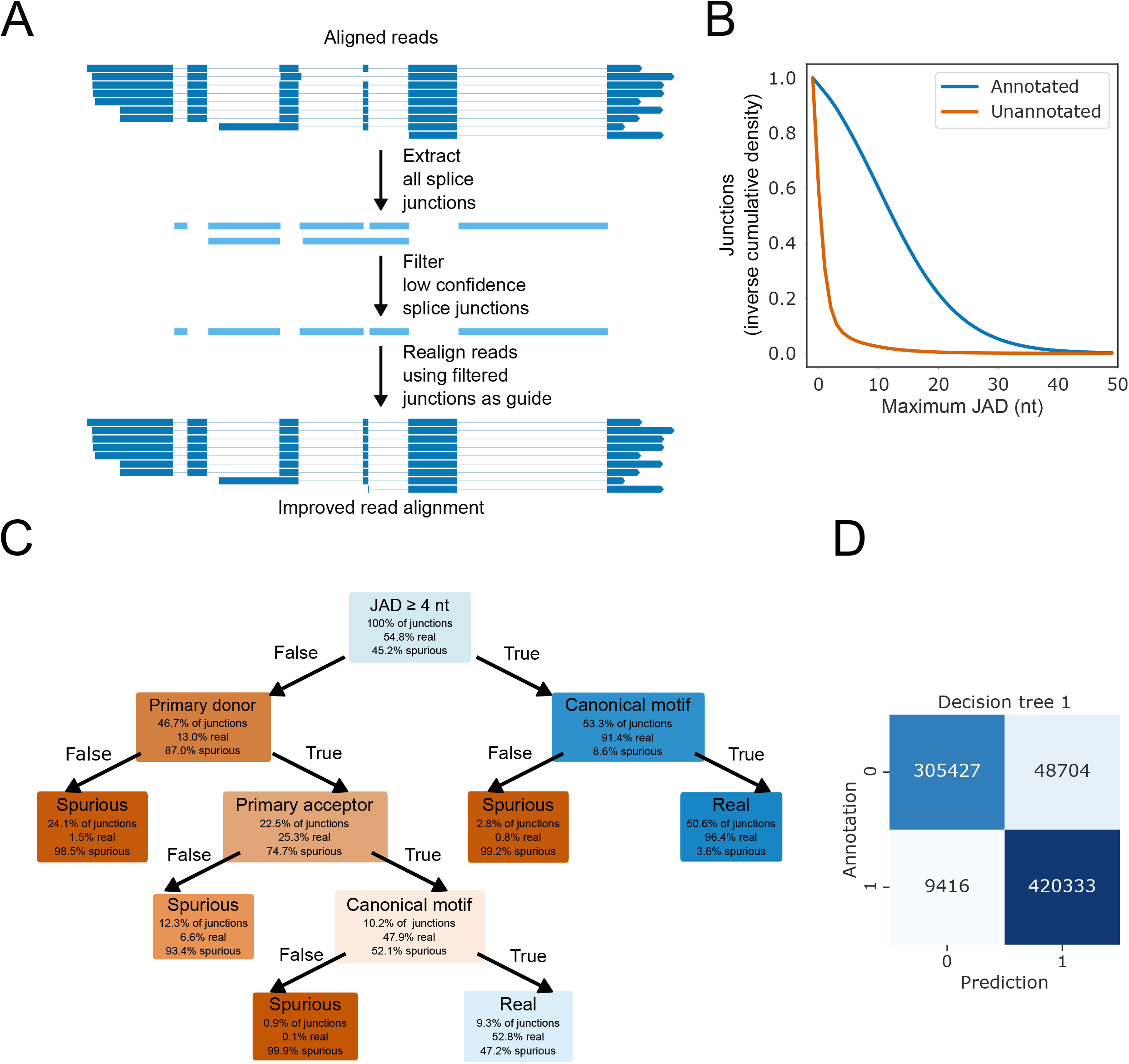
Junction metrics can identify genuine splice junctions. **A** Outline of the two-pass method. **B** The JAD metric can discriminate between annotated and unannotated splice junctions in simulated nanopore DRS reads. Inverse cumulative density plot showing the distribution of per-splice junction maximum JAD values for annotated (blue) and unannotated (orange) splice junctions. **C** Flowchart visualisation of the first decision tree model. Nodes (decisions) and leaves (outcomes) are coloured based on the relative ratio of real and spurious splice junctions. **D** Confusion matrix showing the ratios of correct and incorrect predictions of the first decision tree model on splice junctions extracted from simulated Arabidopsis read alignments.

The junction alignment distance (JAD) is defined as the minimum distance to the first mismatch, insertion or deletion on either overhang of a read alignment splice junction. This metric is used by both finesplice and portcullis software tools(21, 22). For the simulated nanopore DRS read alignment datasets sequenced from Arabidopsis RNA, we found that 88.9% of splice junctions found in the reference annotation had at least one read alignment with a JAD of 4 nt, compared with only 10.1% of unannotated splice junctions (Fig. 3B). Consequently, using a threshold of at least one read with a JAD of 4 nt, we could identify annotated splice junctions with an F1 score of 0.902 (Fig. S3A). Despite the high probability of at least some genuine unannotated splice junctions in the real Arabidopsis data(11), we found that the same JAD threshold could discriminate between annotated and unannotated splice junctions in real datasets to a similar degree (F1 score = 0.899). Similar results were also seen for simulated human datasets, where the same JAD threshold could discriminate between spurious unannotated and genuine annotated splice junctions (F1 score = 0.868). We conclude that the JAD metric is a powerful discriminator of genuine splice junctions across nanopore DRS datasets from different organisms.

Of the other metrics we tested, the read count was predictive of genuine splice junctions at a threshold of >1 read (F1 score = 0.833; Fig. S3B). However, read count correlated strongly with the JAD (Spearman’s ρ = 0.776), suggesting that it does not provide more information. The presence/absence of a canonical intron motif (i.e. GU/AG, GC/AG or AU/AG) had a very high recall, as 99.96% of annotated introns in the simulated alignments were canonical (Fig. S3C). However, the precision was poorer (F1 score = 0.783). This is because in spliced alignment mode minimap2 prefers GU/AG motifs, meaning that 67.1% of spurious splice junctions are also aligned so as to use canonical motifs.

Finally, we developed a primary donor/acceptor metric similar to the one used in portcullis(22). This is calculated by identifying alternative donor or acceptor sites in a 20 nt window around each donor/acceptor and then determining whether they have greater read support than the current site. In case of a tie for read support (e.g. if all splice junctions have a read count of 1), the JAD is used to break the tie, i.e. sites with the largest maximum per-read JAD are considered most likely to be genuine and labelled as a primary site. We found that the primary donor and acceptor metrics were also predictive of genuine splice junctions (F1 scores = 0.842 and 0.785 respectively). By combining the metrics to select splice junctions which are both primary donors and acceptors, the F1 score can be increased to 0.918 (Fig. S3D). It is unclear why the primary donor score is more predictive than the primary acceptor score. A possible reason is that minimap2 is more likely to produce alignment errors at the donor site of splice junctions (e.g. in the case of failure to align small internal exons) or that there are more genuine alternative acceptor sites than donor sites.

We chose to use the identified metrics to create a decision tree model, because these models are easy to interpret and can be kept simple (or pruned) to prevent overfitting. A five-node tree using the JAD, primary donor/acceptor and canonical intron motif metrics (Fig. 3C) was best able to predict genuine Arabidopsis splice junctions (F1 score = 0.935; Fig. 3D). The same decision tree also performed well in predicting genuine and spurious splice junctions from simulated human reads (F1 score = 0.934). This indicates that the model might generalise across nanopore DRS datasets from different organisms, despite their differences in splicing complexity.

### A combination of splice junction alignment metrics and sequence information improves authentic splice junction identification

Genuine splice junctions have sequence biases which are defined by their interactions with spliceosomal uridylate-rich small nuclear RNAs(27). We next asked whether machine learning models could identify genuine splice junctions from the flanking genomic sequences alone. For example, genome sequence information might help identify genuine splice junctions with low read alignment coverage that fail to pass the JAD filter due to stochastic sequencing errors. We therefore extracted 128 nt sequences centred on unique donor and acceptor sites and used these to train LR or random forest models with labels generated by the first decision tree model (Fig. 4A). Using 6-fold cross-validation, we were able to train six models on 83.3% of the data each and use them to make predictions for the remaining 16.7%. Using this approach, we could generate predictions for all splice junctions, with no junction being used for both training and prediction from the same model. We found that LR and random forests performed similarly on the data, indicating that there are few important higher-order interactions (i.e. correlated sequence positions). We therefore proceeded with LR models.

**Fig. 4.**
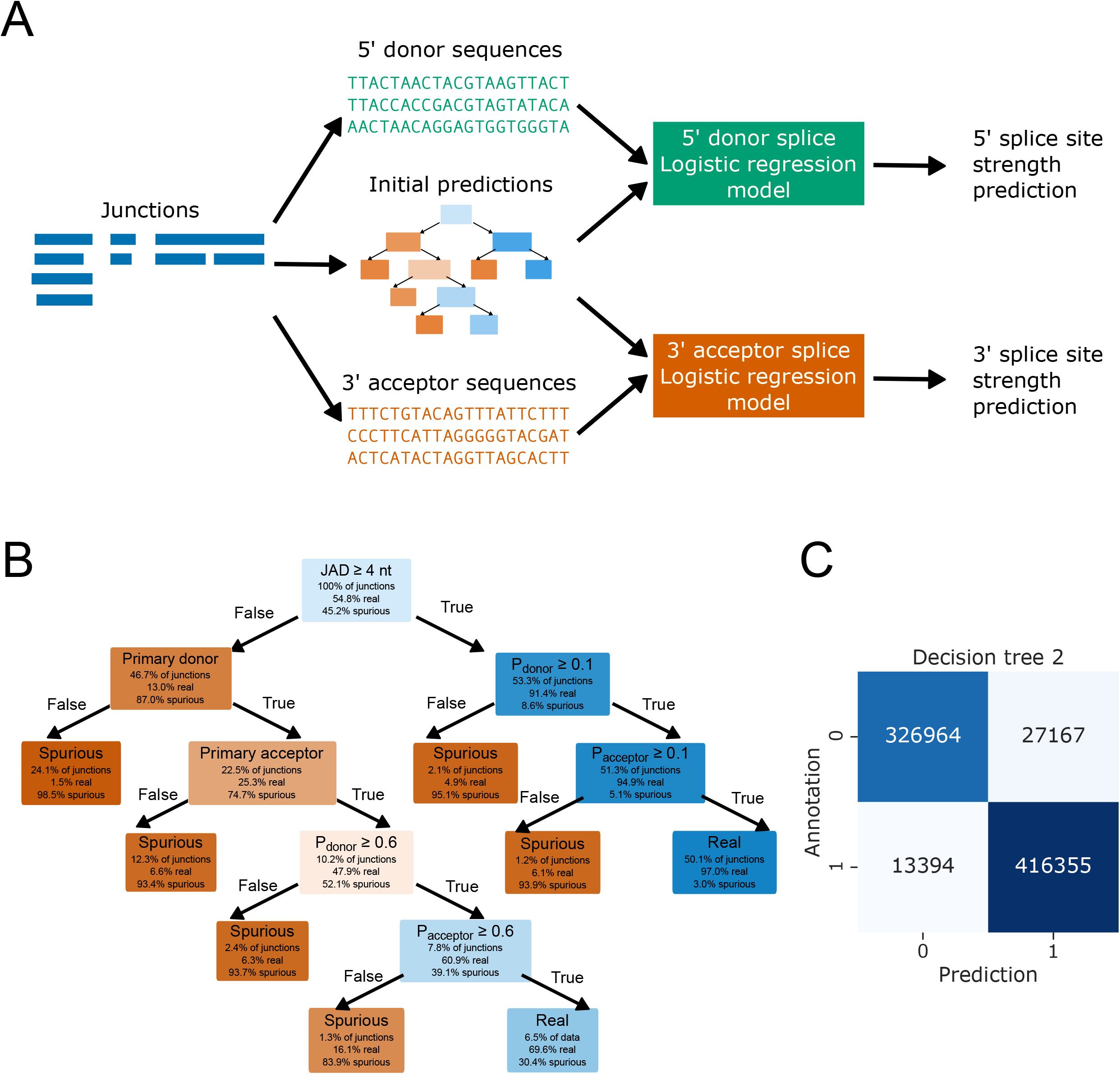
Machine learned sequence information improves identification of genuine splice junctions. **A** Outline of the LR model training process. Sequences from splice junctions were extracted from the reference genome and used as training data (i.e. explanatory variables). Training labels (i.e. the response variable) were generated by the first decision tree model. Independent models were trained for 5′ donor and 3′ acceptor sites and cross-validation used to generate out-of-bag predictions for all sites. **B** Flowchart visualisation of the second decision tree model. Nodes (decisions) and leaves (outcomes) are coloured based on the relative ratio of real and spurious splice junctions. **C** Confusion matrix showing the ratios of correct and incorrect predictions of the second decision tree model on splice junctions extracted from simulated Arabidopsis read alignments.

At a prediction threshold of 0.5, the LR model overclassified positive splice junctions. False positives may be sequences which could in principle act as splice junctions but do not in reality due to effects that the model cannot capture. One such effect could be the presence of alternative splice junctions which are preferentially processed. This is thought to occur under the “first-come-first-served” model of co-transcriptional splicing(28, 29). The model is also unlikely to be able to correctly identify the intron branchpoint motif because this can vary in position relative to the acceptor site(30). Nevertheless, we found that the LR model approach could predict genuine splice junctions from sequence data alone with comparable accuracy to the metric-based decision tree (Fig S4A-C). For example, for the simulated Arabidopsis datasets, using LR on donor and acceptor sequences (with a prediction threshold of 0.5) yielded an F1 score of 0.904 (Fig S4C), which was similar to the F1 score obtained with the JAD or primary donor/acceptor metrics.

We next tested whether the information from the junction metrics and reference sequence model was complementary, i.e. if a combination of the two approaches could produce an improvement in splice junction prediction over each individual approach. Use of a second decision tree model, this time including the JAD metric, primary donor/acceptor metrics and new LR prediction scores (Fig. 4B), further increased the F1 score on splice junctions identified from simulated Arabidopsis read alignments to 0.954 (Fig. 4C). For splice junctions from simulated human reads, we also saw an increase in the F1 score to 0.957. We conclude that an ensemble approach incorporating both junction metrics and sequence information works best for detecting and filtering spurious splice junctions from alignments.

### Two-pass alignment with filtered splice junctions improves transcript identification

We next applied the two decision tree filtering methods to perform two-pass alignment of the simulated reads with minimap2(18). As a positive control, we compared the results to reference-guided alignment with minimap2, since this represents the best possible performance that could be achieved by two-pass alignment (i.e. if the filtered splice junction set perfectly matched the reference annotation). Using filtered splice junctions, the percentage of junctions identified in second-pass alignments that matched annotated splice junctions could be increased over first-pass alignment and naïve two-pass alignment (Fig. 5A). For example, using the simulated Arabidopsis datasets, the median percentage of read alignments matching the splice junctions of the reference transcript they were simulated from increased from 73.2% in the first pass, to 88.2% and 89.3% in a second pass, using the first and second decision tree methods respectively (Fig. 5A). Two-pass alignment rescued the large misalignments of exon 6 seen at *FLM* (Fig. S5A): overall, 86.8% of simulated *FLM* reads aligned to the correct reference transcript after filtered two-pass alignment compared with 19.3% for first-pass alignments. A global improvement in correct alignment was also seen in the simulated human datasets: from 44.4% in the first pass to 64.3% and 65.7% for the two decision tree methods, respectively (Fig. 5A).

**Fig. 5.**
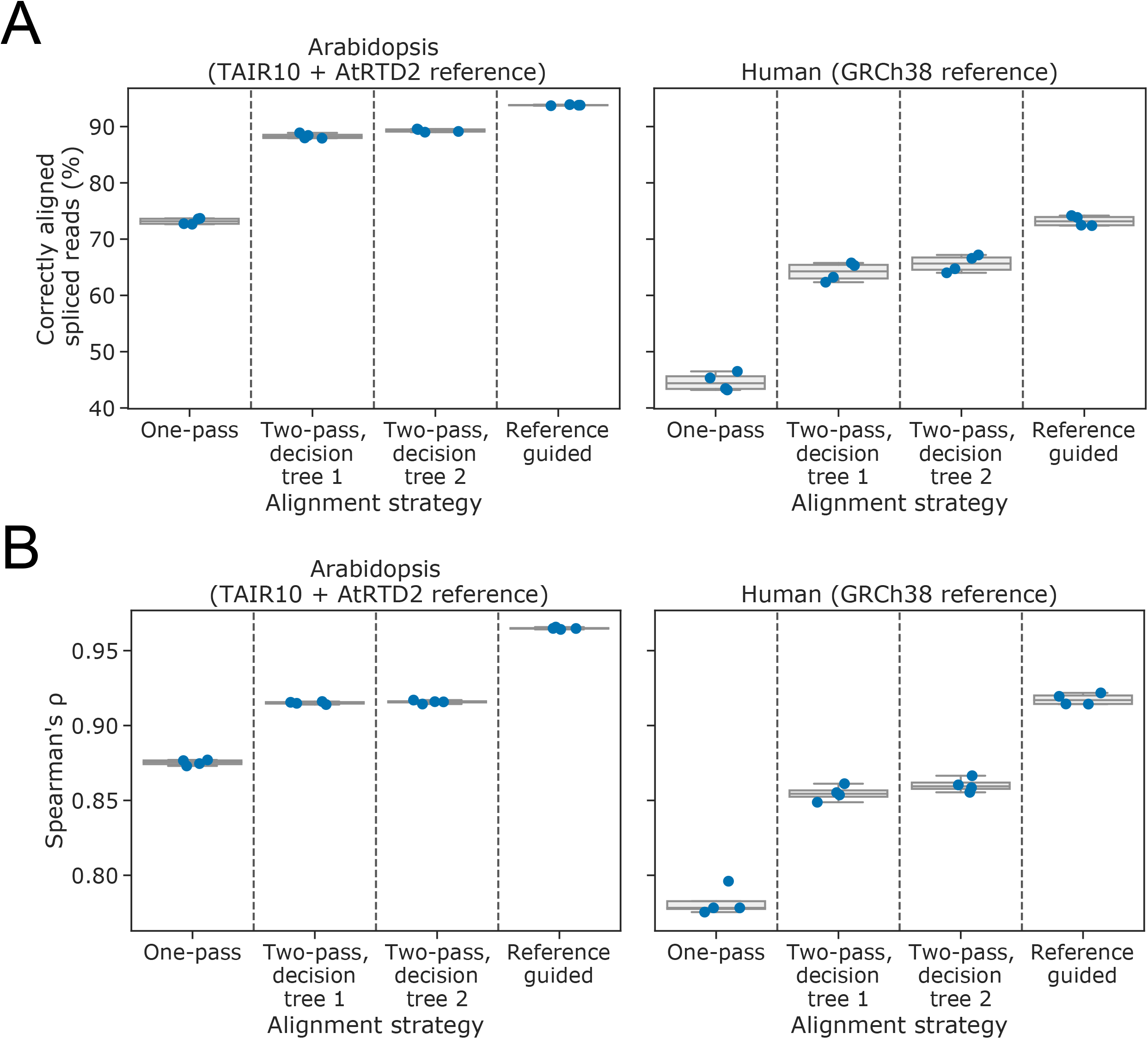
Filtered two-pass alignment improves the identification and quantification of correct transcripts without a reference annotation. **A** Boxplots with overlaid strip-plots showing the percentage of alignments which map exactly to the splice junctions of the transcript from which they were simulated, for one-pass unguided minimap2 alignments, two-pass alignments using splice junctions filtered by decision trees one and two, and reference-annotation-guided minimap2 alignments. Reads were simulated from Arabidopsis TAIR10 + AtRTD2 (left) and human GRCh28 (right) nanopore DRS data. **B** Boxplots with overlaid strip-plots showing the Spearman’s correlation coefficient for actual transcript level counts from simulated data against counts produced by the alignment methods described in A. Reads were simulated from Arabidopsis (left) and human (right) nanopore DRS data aligned to the AtRTD2 and GRCh38 reference transcriptomes, respectively.

Although two-pass alignment improved the number of reads aligning to the correct transcript model, we also detected a slight increase in the number of reads aligning to the wrong annotated transcript. In the simulated Arabidopsis reads analysis, reads aligned using the second decision tree model performed worst on this metric: 2.74% of reads aligned to the wrong isoform compared with only 1.79% of reads after first-pass alignment (Fig. S5B). To assess whether such misassignment affects the quantitation of transcripts, we calculated Spearman’s correlation coefficient (ρ) for estimated versus known transcript level read counts for the simulated data (Fig. 5B). The results indicated that, despite this misassignment, two-pass aligned reads could be quantified accurately, with an overall improvement in median Spearman’s ρ for one-pass versus two-pass of from 0.876 to 0.916 for simulated Arabidopsis reads (Fig. 5B) and from 0.778 to 0.859 for simulated human reads (Fig. 5B). However, there may be corner cases where transcript misassignment could have consequences for transcript usage analysis. This should be considered for experiments where quantification is important. Overall, we conclude that two-pass alignment using filtered junctions can improve both the detection of correct splicing patterns and the quantitation of nanopore DRS reads.

### Filtered two-pass alignment improves reference-guided annotation

Summarising read alignments into annotations facilitates transcript level quantification of short and long reads and aids the interpretation of RNA processing complexity. We therefore asked whether two-pass alignment of spliced long reads with relatively high sequence error rates can improve the results of genome-guided annotation tools. Several software tools designed to produce annotations from long reads exist, including FLAIR(10) and pinfish (ONT), which were designed for nanopore DRS data; TAMA(31), which was designed for PacBio IsoSeq data; and StringTie2(6), which was designed as a technology-agnostic long-read assembly tool.

We benchmarked our methods using StringTie2 because it is reported to be faster and more accurate than FLAIR on simulated nanopore DRS data(6). Using full-length reads simulated from real Arabidopsis and human nanopore DRS data, we could identify the intron-chain-level precision and recall of annotations assembled from reads processed using either one-pass or two-pass alignment. Here, precision is defined as the percentage of assembled transcripts whose combination of introns match a transcript in the reference annotation; and recall is defined as the percentage of annotated transcripts for which at least one read was simulated and whose combination of introns match a transcript assembled from simulated reads. We assessed reads aligned using guide splice junctions from the reference annotation as a positive control.

For both Arabidopsis and human datasets, two-pass alignment generally produced a clear improvement in both precision and recall of StringTie2 transcript assembly over first-pass alignment (Fig. 6A). Of the two decision tree methods produced, decision tree 2 (using junction sequence information) performed best (median F1 score was 0.699 for the Arabidopsis data and 0.629 for the human data). There was a particularly large increase in precision for reference annotation-guided alignment of at least 8.7% and 9.6% over one-pass alignment for all Arabidopsis and human samples, respectively (Fig. 6A).

**Fig. 6.**
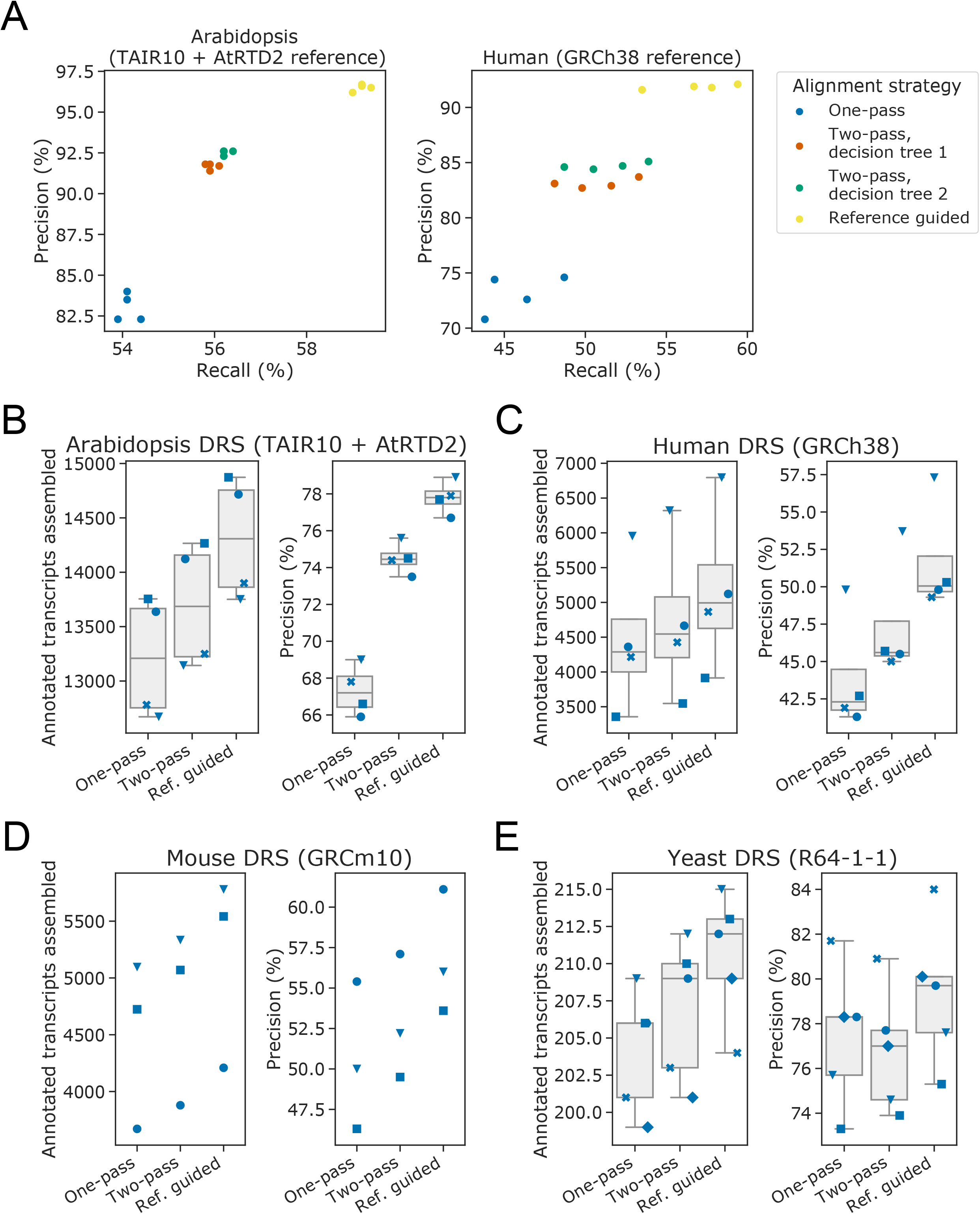
Filtered two-pass alignment improves genome-guided annotation. **A** Scatterplot showing precision against recall for intron chains in genome-guided transcriptome annotations generated from alignments using StringTie2. Precision and recall scores were calculated against reference annotations filtered to include only transcripts for which at least one read was simulated. Reads were simulated from Arabidopsis (left) and human (right) nanopore DRS data aligned to the AtRTD2 and GRCh38 reference transcriptomes, respectively. **B–E** Stripplots with box-and-whiskers showing the number of correct transcripts assembled (left panels) and precision of transcripts assembled (right panels) for genome-guided transcriptome assembly using StringTie2. Two-pass alignment improved the precision and number of transcripts assembled for real nanopore DRS data for **B** Arabidopsis, **C** human, **D** mouse and **E** yeast. For all boxplots, overlaid strip-plots are shown for individual samples. Each sample was assigned a unique marker so that the changes in each sample could be tracked between the one-pass, two-pass and reference-guided alignments. Box-and-whiskers not shown for samples with less than 4 data points. Y limits vary between figures since within-figure (i.e. same species and sequencing technology) comparison is more important than between-figure comparisons.

We next considered whether two-pass alignment could improve the genome-guided transcriptome assembly performance of Stringtie2 on real datasets, using current reference annotations as a ground truth. However, it is important to note that there may be genuine transcript examples in the datasets that are not yet included in the reference annotation; if so, this will affect the measurement of precision. Furthermore, recall against the reference is likely to depend on the sequencing depth of samples. We therefore report the number of annotated transcripts assembled for each sample, rather than the recall.

Two-pass alignment improved both the precision and the number of transcripts assembled for Arabidopsis, human and mouse samples(10, 11, 32) (Fig. 6B–D). This approach resulted in a median increase in assembly precision compared with one-pass alignment of 7.1% for Arabidopsis samples, 3.5% for human samples and 2.2% for mouse samples (median increase in annotated transcripts assembled per sample of 478.5, 257.5 and 238, respectively). We conclude that for organisms with complex patterns of pre-mRNA splicing, two-pass alignment can improve both the precision and number of correct (annotated) transcripts assembled by StringTie2 from real nanopore DRS data.

When we applied the same approach to the yeast *Saccharomyces cerevisiae*, the results were very different (Fig. 6E). In this species, two-pass alignment resulted in a median increase of only three more annotated transcripts assembled per sample and an increased number of unannotated transcripts assembled, resulting in a median decrease of 0.8% in assembly precision. Splicing complexity in *S. cerevisiae* is relatively low: there are only 364 annotated introns in the Ensembl R64 annotation, most genes are intronless, and most introns are constitutive(33). This led to a high ratio of unannotated splice junctions in first-pass alignments (the median number of junctions identified was 8,056), suggesting that the vast majority of junctions in the dataset are spurious. Furthermore, most *S. cerevisiae* introns occur close to mRNA 5′ ends, resulting in typically short upstream exons that present challenges to alignment software. Such a large ratio of spurious to genuine splice junctions is likely to affect the precision of junction filtering. Notably, even when the reference annotation was used to guide alignment, precision was only improved by a median of 1.9% (with a median of six more transcripts assembled correctly). Intron-containing genes are generally more highly expressed (many encode ribosomal proteins) than intronless genes(34). This may mean that the coverage of annotated transcripts is already good and, thus, that the number of true annotated transcripts assembled cannot be much improved. This result suggests that both reference annotation-guided and two-pass alignment methods have limited use for genome-guided transcriptome assembly in organisms with low complexity splicing.

Finally, we considered whether filtered two-pass alignment could improve genome-guided annotation of nanopore DRS reads derived from sequencing cDNA copies and from PacBio IsoSeq data (Fig S6A-D). To assess this, we used the recommended alignment parameters for minimap2(18), but with the splice junction filtering parameters that were used for nanopore DRS data. Overall, the precision and recall of transcripts assembled from both nanopore cDNA and PacBio IsoSeq data for human, mouse and Arabidopsis samples could be improved using two-pass alignment. For human and mouse nanopore cDNA samples, two-pass alignment resulted in a median increase of 3.85% and 2.3% in assembly precision, respectively, compared with one-pass alignment (median increase in annotated transcripts assembled per sample of 609.5 and 420.0, respectively; Fig. S6A,B). For Arabidopsis and human PacBio IsoSeq samples, two-pass alignment resulted in a median increase of 8.45% and 1.35% in assembly precision, respectively, compared with one-pass alignment (median increase in annotated transcripts assembled per sample of 63 and 242.5, respectively; Fig. S6C,D). We conclude that a two-pass method can improve genome-guided transcript assembly of the high-error long reads produced using a range of sequencing technologies.

### Two-pass alignment can aid novel splice-isoform discovery in annotated species

We have shown that a two-pass approach can improve the accuracy of spliced alignment in the absence of a reference annotation. However, even the most well-studied genomes are likely to be incompletely annotated, and so novel splice-junction discovery which builds upon existing annotations is also desirable. We therefore developed an alternative two-pass method which allows users to provide reference annotations. The annotation is used to train random forest models which can then predict novel splice junctions. These models replace the pre-trained decision trees used in the annotation-independent method. We refer to this method hereafter as annotation-aided two-pass alignment.

If a reference annotation for a species is truly complete – i.e. there are no new splice-junctions to be discovered, then two-pass alignment can only reduce the accuracy of alignment by introducing false-positive introns into the guide splice junction set. We therefore hypothesise that two-pass alignment will be useful when many genuine splice junctions are missing from the annotation, because genuine novel splice junctions added to the guide junction set will outweigh false-positives that are introduced. We refer to the percentage of genuine splice junctions that are unannotated as the level of annotation “missingness”. To test our hypothesis, we performed random subsampling of transcript isoforms in the Arabidopsis reference annotation to simulate an incomplete reference at a range of missingness levels, from 0.1% to 90% missing. We then performed annotation-aided two-pass alignment of the nanopore DRS dataset and assessed the predictive performance on splice junctions which were absent from the subsampled annotation. We found that the annotation-aided method performed best for medium missingness levels. For example, in Arabidopsis DRS data, when between 25% and 66% of reference isoforms were missing, the true positive rate / recall was high (minimum of 0.86), for a low false positive rate (maximum 0.15) and a high precision (minimum 0.85) (Figure 7A,B). This translates to a 1.3-3.9% improvement in the percentage of correctly aligned reads compared to reference-guided alignment (Figure 7C). At missingness levels of less than 25%, the false positive rate increased and precision decreased (Figure 7A,B). The reason for this decrease in performance is because as the reference annotation nears completion, the imbalance between genuine novel splice junctions and false positives caused by alignment errors increases. However, reductions in splice-junction level precision do not translate to a large drop in the percentage of correctly aligned reads – at 0.1% missingness, the reduction was 0.36% (Figure 7C). Furthermore, at lower levels of missingness, the recall remained high, with at least 96.7% of all genuine novel splice junctions being detected. At extremely high levels of annotation missingness, the recall of the two-pass filtering method begins to fall – at 90% missing, recall is only 0.12 (Figure 7B). This is likely to be because when the reference is extremely incomplete, it no longer represents a good training dataset, since a large proportion of junctions missing from the reference will be genuine. For reference missingness levels >75%, it was therefore better to perform two-pass alignment without the reference annotation (Figure 7C,D). With human RNA datasets, we found that annotation-aided two-pass alignment improved the percentage of correctly aligned reads when transcript isoform missingness was at least 25% (Figure 7D). This is likely due to the completeness of human annotation – more junctions are found in more than one transcript isoform. We conclude that annotation-aided two-pass alignment is most useful when a high-quality annotation is available, but where the conditions of the experiment are expected to produce a significant number of novel splice junctions.

**Fig. 7.**
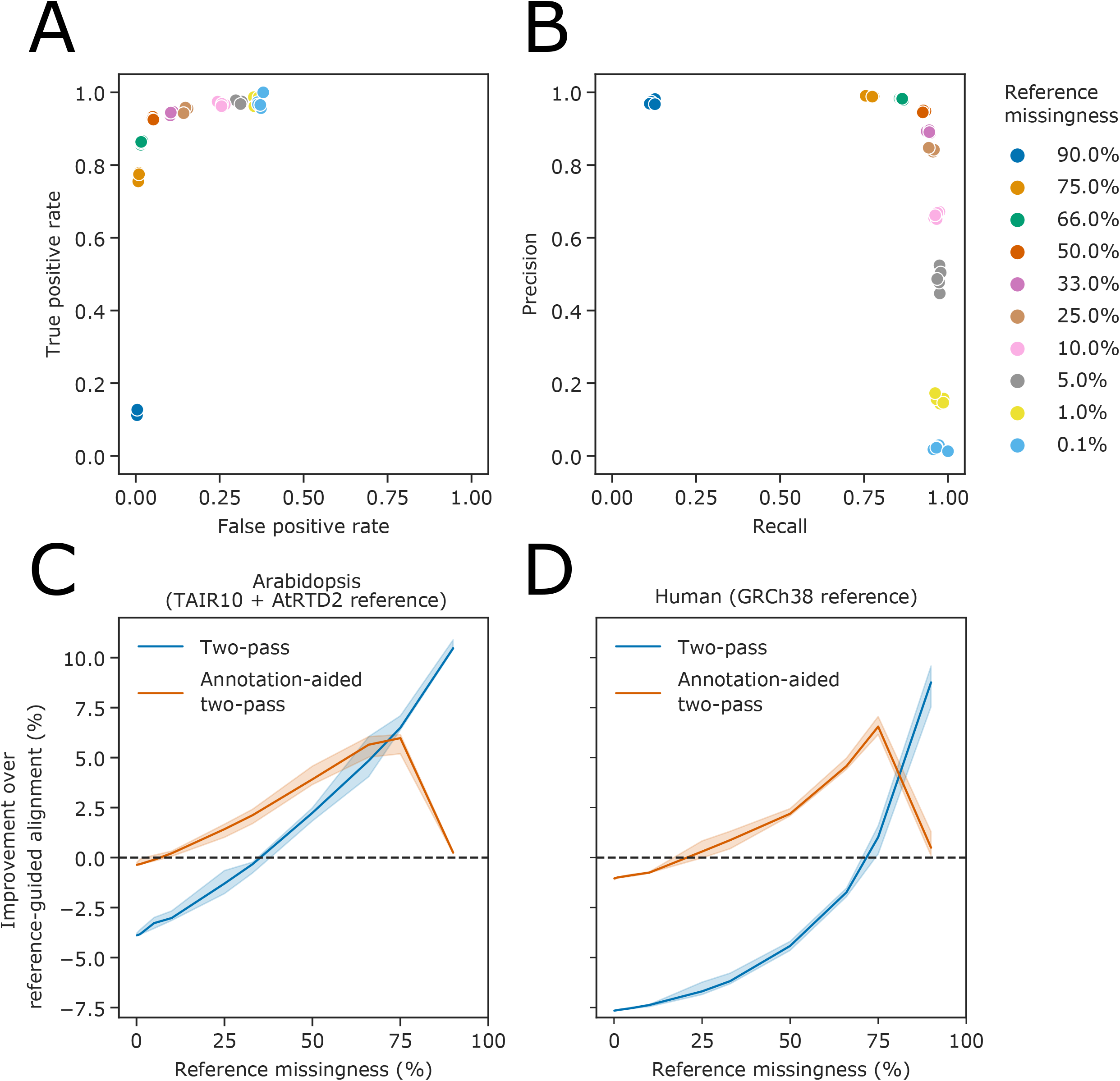
Annotation-aided two-pass alignment rescues missing splice junctions. **A** ROC scatterplot and **B** precision/recall scatterplot showing true positive rate and false positive rate of novel splice junction classification in simulated Arabidopsis read alignments, at different rates reference annotation missingness. Annotated transcript isoforms were subsampled to simulate incomplete reference annotations, and these were used to inform annotation-aided two-pass alignment. **C-D** Line plots showing the improvement in the percentage of correctly aligned reads using two-pass alignment compared to reference-guided alignment at different reference annotation missingness rates for **C** Arabidopsis and **D** humans, respectively. Blue line shows improvement compared to reads aligned using two-pass method only. Orange line shows improvement compared to reads aligned using reference-annotation in first-pass, followed by annotation-aided junction filtering and second pass alignment. Shaded regions represent 95% confidence intervals.

### Two-pass alignment discovers novel splice isoforms in the Arabidopsis RNA exosome mutant hen2-2

To validate the annotation-aided two-pass approach, we performed a case study with Arabidopsis using the *hen2-2* mutant. HEN2 functions as an accessory protein to the nuclear RNA exosome, and is required for the processing and degradation of specific classes of mRNAs and non-coding RNAs(35). As a result, many RNAs, some of which contain novel splice junctions, accumulate in the *hen2-2* mutant compared to wild-type. Many of these transcripts are unannotated because exosome mediated decay means that they are effectively “hidden” in wild-type plants. We have previously performed Illumina RNAseq of *hen2-2* mutants at relatively high depth(11). We therefore generated nanopore DRS reads from similar tissue and performed annotation-aided two-pass alignment to detect novel splice junctions. Of the 17,521 unannotated splice junctions detected in first-pass alignment of the nanopore DRS data, only 20% (3548) are supported by Illumina RNAseq, and only 24% (4210) passed filtering, indicating that the majority are spurious (Figure 8A). However, of those that pass filtering, 57% (2382) were supported by Illumina RNAseq. This represents 67% of the 3548 unannotated junctions which were supported by both nanopore DRS and Illumina RNAseq. For example, we detected novel isoforms of annotated genes, such as *AT1G19396*, where use of an alternative donor site in a large intron results in a novel exonic region (Figure 8B). We also detected completely unannotated transcripts, such as an antisense RNA at *AT3G12140* with multiple novel splicing events (Figure 8C). We conclude that two-pass alignment is able to detect genuine novel introns in well-annotated species, under less well-annotated conditions.

**Fig. 8.**
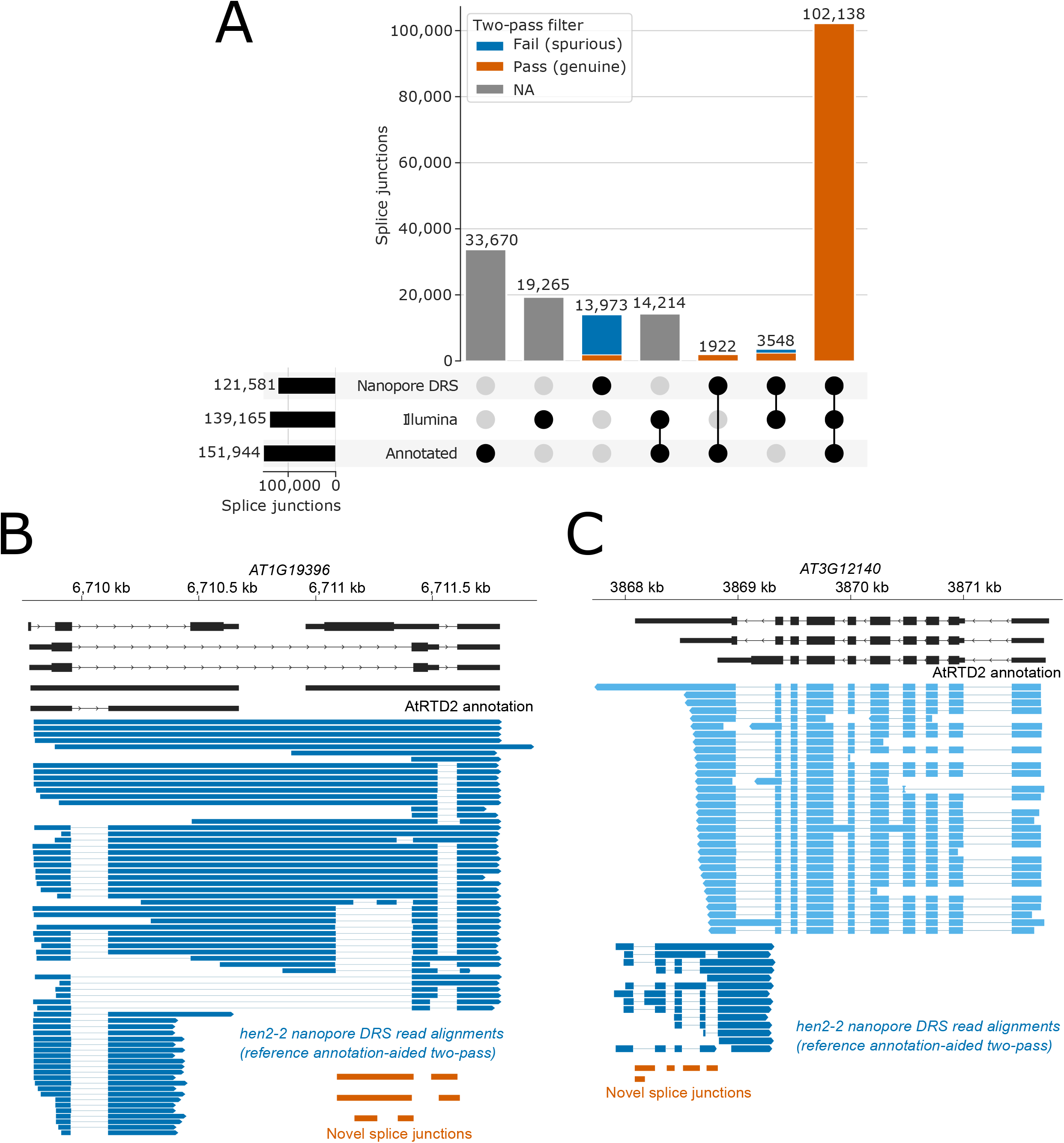
Annotation-aided two-pass alignment identifies novel splice isoforms in *hen2-2* mutants. **A** Upset plot showing the intersection of splice junctions detected using nanopore DRS or Illumina RNAseq, and presence in the AtRTD2 annotation. Horizontal bars show the overall number of junctions detected using each technology/annotation, whilst stacked vertical bars represent set intersections. For nanopore DRS data, splice junctions with one or more supporting read alignment are shown. For Illumina RNAseq, splice junctions with ten or more supporting read alignments are shown. Nanopore DRS junctions which are classified as spurious by the two-pass filtering method are labelled in blue, whilst junctions which are classified as genuine are labelled in orange. Set intersection bars not including nanopore DRS are shown in grey. **B-C** Gene track showing novel splice isoforms detected at **B** *AT1G19396* and **C** *AT3G12140* in *hen2-2* nanopore DRS data. AtRTD2 annotation is shown in black. Nanopore DRS reads are shown in blue (positive strand) or light blue (negative strand). Novel splice junctions are shown in orange.

### Conclusions

RNA sequencing is a fundamental tool for understanding what genomes really encode. Technological approaches that directly sequence full-length RNA molecules substantially increase the useful information that RNA sequencing can provide. The challenges that alternative splicing, in particular, presents to the interpretation of high-throughput RNA sequencing data means that software development needs to accompany progress in sequencing technology. In this way, knowledge gained from ambitious genome sequencing programmes such as the Earth BioGenome Project, which aims to characterise all eukaryotic life on Earth(36), can be maximised. We have shown that a two-pass alignment approach, informed by splice junction alignment metrics and machine learning of sequence features associated with splicing, can improve the accuracy of intron detection in long-read data. Knowledge of existing splice junctions can also be applied to aid the discovery of novel splicing events when annotations are incomplete - for example, in disease states with altered gene expression. Consequently, this approach can enhance the utility and realise the potential of long-read RNA sequencing.

## Methods

### Nanopore and PacBio data

Four replicates of nanopore DRS reads derived from Arabidopsis Col-0 RNA were used (11). These datasets are available in FAST5 format from the European Nucleotide Archive under accession no. PRJEB32782. The first four listed replicates of DRS and cDNA sequencing reads derived from human cell line GM12878 were used: Birmingham DRS samples 1, 2, 3 and 5; Birmingham cDNA samples 1 and 2; Hopkins cDNA samples 1 and 2)(10). DRS datasets were downloaded in FAST5 format and cDNA datasets in FASTQ format using the links provided on GitHub (http://s3.amazonaws.com/nanopore-human-wgs/rna/links/NA12878-DirectRNA_All.files.txt). Mouse DRS and cDNA datasets in FASTQ format (32) were downloaded from the European Nucleotide Archive (accession no. PRJEB27590). Yeast DRS datasets in FASTQ format (9) were downloaded from the European Nucleotide Archive (accession no. PRJNA408327). Human IsoSeq datasets in FASTQ format were downloaded from the PacBio AWS webserver (http://datasets.pacb.com.s3.amazonaws.com/2014/Iso-seq_Human_Tissues/list.html). Arabidopsis IsoSeq data in FASTQ format was downloaded from the European Nucleotide Archive (accession no. PRJNA371677).

### hen2-2 nanopore DRS data

For newly sequenced nanopore DRS data, *hen2-2* seeds were sown on MS10 medium plates, stratified at 4°C for 2 days, germinated in a controlled environment at 22°C under 16 hr light/8 hr dark conditions and harvested 14 days after transfer to 22°C. RNA isolation and nanopore direct RNA sequencing were performed as described previously(11).

### Preliminary data processing

Pipelines for processing of data were written using snakemake version 5.10.0(37). FAST5 data was re-basecalled locally using guppy version 2.3.1 (ONT). All alignments were performed using minimap2 version 2.17-r963(18). Arabidopsis reads were aligned to the TAIR10 reference genome(38) and AtRTD2 reference transcriptome(23). Human, mouse and yeast reads were aligned to the GRCh38, GRCm38 and R64–1-1 primary assemblies and to cDNA transcriptomes from Ensembl, respectively(24). Alignments to reference genomes were performed using spliced parameters. For DRS datasets, these were: -k14 -x splice -L --cs=long. For nanopore cDNA and PacBio datasets the parameters used were -x splice -L -- cs=long. The maximum intron size (-G) was set at 10,000 nt for Arabidopsis samples, at 200,000 nt for human and mouse datasets and at 5,000 nt for yeast, to match the known intron length distributions in these organisms. For two-pass alignments using a guide splice junction set, a junction bonus (--junc-bonus) of 12 was also used, as this was found to improve the percentage of correctly aligned simulated reads when performing reference-guided annotation, compared to the default (--junc-bonus 9). Alignments of DRS reads to the reference transcriptome were performed using splicing-free parameters, namely: -k14 --for-only -L --cs=long.

### Simulation of DRS reads

To provide a ground truth with a complete set of known splice sites, sequences were simulated from the reference transcriptomes, with length and error profiles matching those of real DRS reads. This was done by modelling the length, homopolymer error and other error profiles of real reads. Only primary alignments were considered. The cs tags of reads aligned to the reference transcriptome were used to recreate pairwise alignments between each read and the reference, ignoring refskips. Alignments were inverted to match the 3′ → 5′ sequencing direction of nanopore DRS. Aligned basecalls at reference homopolymers of ≥5 nt in length were used to build a probability model of homopolymer calls given the reference homopolymer. To prevent these error profiles being modelled multiple times, the reference homopolymer was then replaced with the aligned basecall in the pairwise alignment. Next, the altered alignment was used to create a Markov chain model of basecalled sequence given the reference sequence. For each base in the reference sequence in the alignment, the aligned portion of the query sequence was identified. The “state” of the alignment (i.e. match, mismatch, insertion or deletion) was also identified. The probability of seeing a query sequence was calculated, given the current and previous four bases of the reference and the previous four states of the alignment.

The reference transcriptome was also used to simulate data using these models. The number of primary alignments in the real data for each reference transcript was used as the number of simulated reads per transcript. To simulate basecall errors, sequences were inverted to the 3′ → 5′ direction and reads were generated using Markov chain Monte Carlo simulations with the basecall model. The reference sequences were prepended with a 10 nt oligo(A) sequence to mimic a short poly(A) tail so that the initial state of the Markov chain was always “AAAAA” and “====” (i.e. four matches). Homopolymers in the simulated read were identified and replaced with randomly selected sequences from the homopolymer model. The read was then reverted to the 5′ → 3′ direction for mapping. Because we wanted to assess the alignment of full-length reads, we did not model or simulate the 3′ bias, which is inherent to nanopore DRS data. However, 10 nt of simulated read were subtracted from the 5′ end of reads to simulate loss of signal at the end of sequencing.

### Post-alignment splice junction correction with FLAIR

BAM files were converted to the BED12 format using bedtools(39). BED12 files were then corrected using the reference GTF annotation with FLAIR correct version 1.4 and default settings(10).

### Junction metric calculations

Splice junctions and junction metrics were extracted from aligned reads using the long form cs tag produced by minimap2 version 2.17(18) using pysam version 0.15.4. The per-read JAD was calculated as the length of the shorter of the two match operations immediately flanking refskip (splicing) operations. Where there were mismatches or indels immediately adjacent to refskips, a JAD of zero was assigned. The per-splice junction JAD was calculated as the maximum of the per-read JADs. Intron motifs were extracted from cs tags. For Arabidopsis, human and mouse samples, GU/AG, GC/AG and AU/AG splice junctions were all considered canonical. For yeast samples, only GU/AG splice junctions were considered canonical. To calculate the primary donor/acceptor metrics, interval trees of donor and acceptor sites were constructed using NCLS(40). Donors were assigned as primary donors if there were no alternative donor sites within 20 nt with higher read counts. Likewise, acceptors were considered primary if there were no alternative acceptors within 20 nt with higher read counts. Ties were broken using the JAD metric, i.e. the splice junctions with higher JADs were assigned primary status. Where there were still ties after read count and JAD comparisons, no splice junctions were assigned primary status. Splice junctions extracted from four replicates of Arabidopsis or human DRS reads were used to build decision tree models with scikit-learn version 0.22.1(41). A minimum depth of 4, minimum number of samples required to split a node of 1000, and minimum Gini impurity decrease required to split a node of 0.005 were used. The decision tree generated from Arabidopsis reads was a subtree of the human tree (i.e. it could be created by pruning the human tree), indicating that the decision function can generalise across samples.

### Reference sequence filtering using LR models

Splice junctions obtained from a first-pass alignment were separated into lists of unique donor sites and unique acceptor sites. These were labelled as positive training examples if they participated in at least one donor/acceptor pair which passed the first decision tree function. Sequences of 128 nt for each splice junction (centred on the donor or acceptor site) were extracted from the reference genome using pysam version 0.15.4 and one hot encoded into four binary variables to create a 512-feature training dataset. LR models were trained using 6-fold cross-validation with scikit-learn version 0.22.1(41). For each fold, the model was used to generate out-of-bag predictions on the held-out data. The probabilities produced were then used in place of the canonical intron motif to produce the second decision tree, using a maximum depth of 6, a minimum number of samples of 1,000 and a minimum Gini impurity decrease of 0.003. Thresholds for splice scores in the tree were simplified to comprise only a high confidence threshold of 0.6 (for rescuing splice junctions failing the JAD metric threshold) and a low confidence threshold of 0.1 (for removing false positives from junctions passing the JAD metric threshold).

### Annotation-aided two-pass alignment

For use cases where high quality annotations are already available, we developed an annotation-aided two-pass approach. Here, annotated junctions are provided along with read alignments. Annotated junctions are labelled as genuine. Unannotated junctions discovered in alignments are assumed to be mainly spurious. These labels are then used to train an extremely random forest model on junction metrics. Out-of-bag predictions for each junction are used as refined labels for LR models to detect splice junction sequence. A final extremely random forest model is trained on refined labels, using junction metrics and splice junction sequence scores determined by LR models. Positive examples which are not in the annotation will be a mixture of false positives and genuine novel splice-junctions. Any false negatives from the annotation are (optionally) retained.

### Evaluation of splice junction models

Performance of the metrics and models was evaluated at splice junction level using the reference annotation as a ground truth. For simulated datasets, annotation is the absolute ground truth because all reads are simulated using only splice junctions in the annotation. For real datasets, some “false positives” are likely to be genuine splice junctions and some junctions in the reference, which appear as false negatives, are actually incorrectly annotated or not expressed. Precision is defined as the number of true positives divided by the total number of positive predictions by the model, i.e. true positives ÷ (true positives + false positives). Recall is defined as the number of true positives divided by the total number of real positive examples in the dataset, i.e. true positives ÷ (true positives + false negatives). The F1 score is the harmonic mean of the precision and recall.

### Evaluation of alignments

To evaluate alignments, we used the intron chain of reference transcripts as a ground truth. The intron chain is the pattern of linked splicing in a transcript, disregarding the transcription start and termination sites. Alignments of simulated reads were considered correct if they mapped correctly to the intron chain of the reference transcript they were simulated from, with no mistakes. Simulated reads that were mapped using intron chains not included in the reference or as being intron-less when they should have splicing were considered novel spurious alignments. Simulated reads that were mapped using the intron chain of a reference transcript other than the transcript they were simulated from were considered to be misassigned. For measures of quantification accuracy, alignment counts for transcripts were generated using the number of simulated reads that aligned with the same splice junctions as the reference transcript. Spearman’s correlation coefficients were then calculated against the known input transcript counts for simulation.

### Reference-guided assembly

Reference-guided transcriptome assemblies were produced using StringTie2(6) version 2.1.1 in long-read mode, with otherwise default parameters.

### Evaluation of assemblies

Reference-guided transcriptome assemblies were evaluated using the precision and recall of intron chains calculated using gffcompare with default settings(42). The input reference GTF files were filtered to include only transcript models for which at least one read had been simulated.

### Reference missingness analysis

To simulate incomplete references, transcript isoforms were removed from the Araport11 (Arabidopsis) and GRCh38 (human) reference annotations at rates from 0.1% to 90%. These incomplete references were then used to perform reference guided alignment of reads simulated using the full reference annotation. Splice junctions from read alignments were then filtered using the annotation-aided method, and reads were realigned using filtered junctions as a guide. Performance on splice-junctions was measured on junctions which were not present in the annotation (i.e. training set) only. Performance at read-alignment level was measured as the change in the percentage of correctly aligned reads compared to using only the incomplete reference annotation to guide alignment.

### Illumina RNAseq analysis

*hen2-2* Illumina RNAseq data was downloaded from PRJEB32782. Reads were mapped to the TAIR10 genome using STAR, with a splice junction database built from the Araport11 annotation. Splice junction set intersections were identified in Python using pysam, and the visualised using upset plots.

## Declarations

### Availability of data and materials

#### Code availability

The methods used to filter splice junctions have been implemented in the “2passtools” python package, which is available on GitHub in repository https://github.com/bartongroup/2passtools. The software used to simulate reads is available on GitHub in repository https://github.com/bartongroup/yanosim.

The scripts, pipelines and notebooks used to perform benchmarking and generate figures are available on GitHub in repository https://github.com/bartongroup/two_pass_alignment_pipeline.

#### Data availability

Basecalled and simulated nanopore DRS datasets are available from Zenodo at https://zenodo.org/record/3773729. Newly generated nanopore DRS FAST5 data has been made available on ENA under accession PRJEB41381.

### Author contributions

MTP developed the software and performed the data analysis. KK performed the *hen2-2* nanopore DRS sequencing. MTP and GGS wrote the manuscript. All authors commented on the manuscript.

## Acknowledgments

We thank James Abbott for testing the pipeline.

## Supplemental Figures

**Fig. S1.**
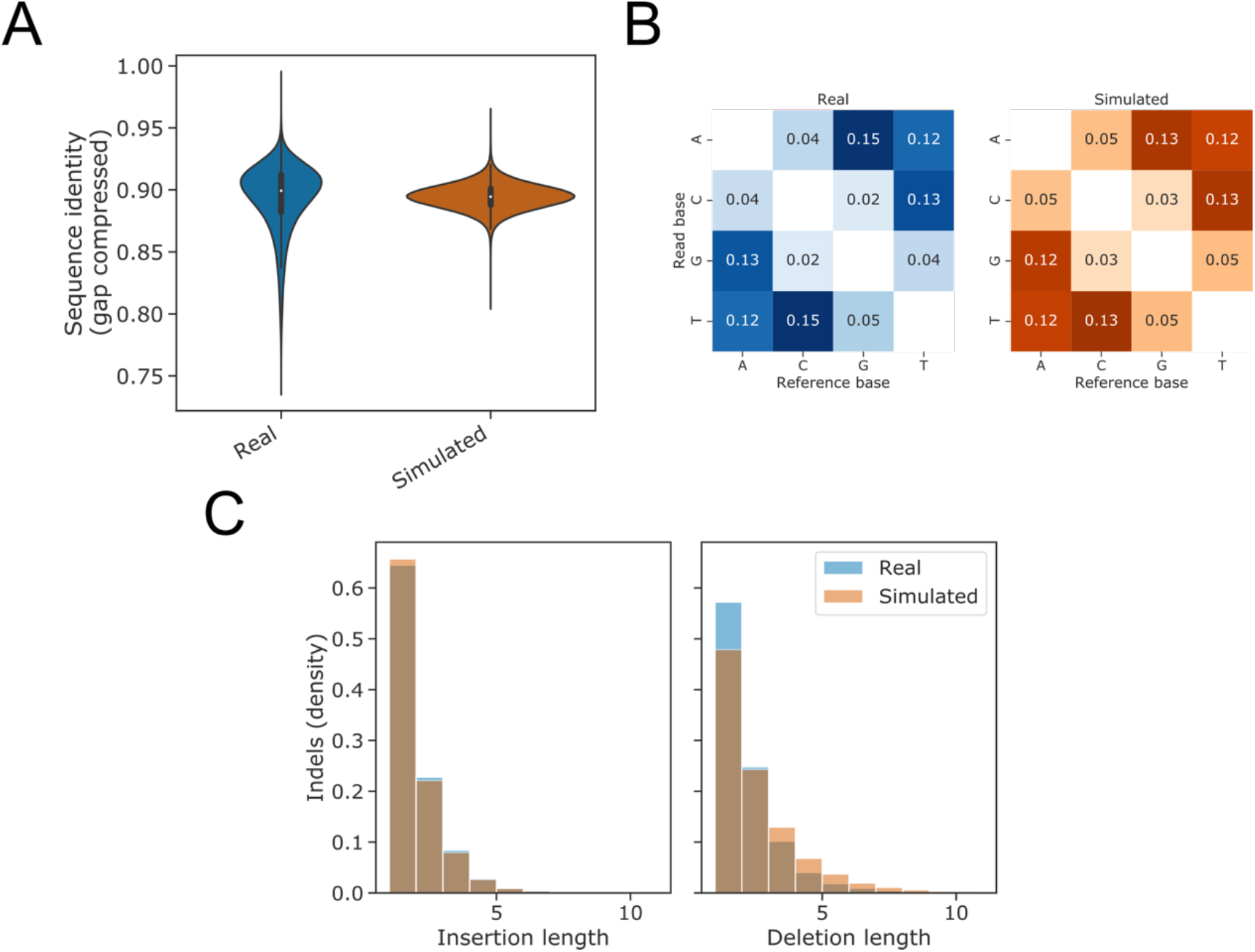
Simulation of nanopore DRS read alignments. **A** Violin plot showing the distribution of sequence identity scores for real and simulated Arabidopsis nanopore DRS reads. Simulated reads match the median sequence identity of real reads, although they do not capture the tails of high- and low-quality reads. **B** Insertion and deletion length distributions for real and simulated nanopore DRS reads. **C** Mismatch profiles for real and simulated nanopore DRS reads.

**Fig. S2.**
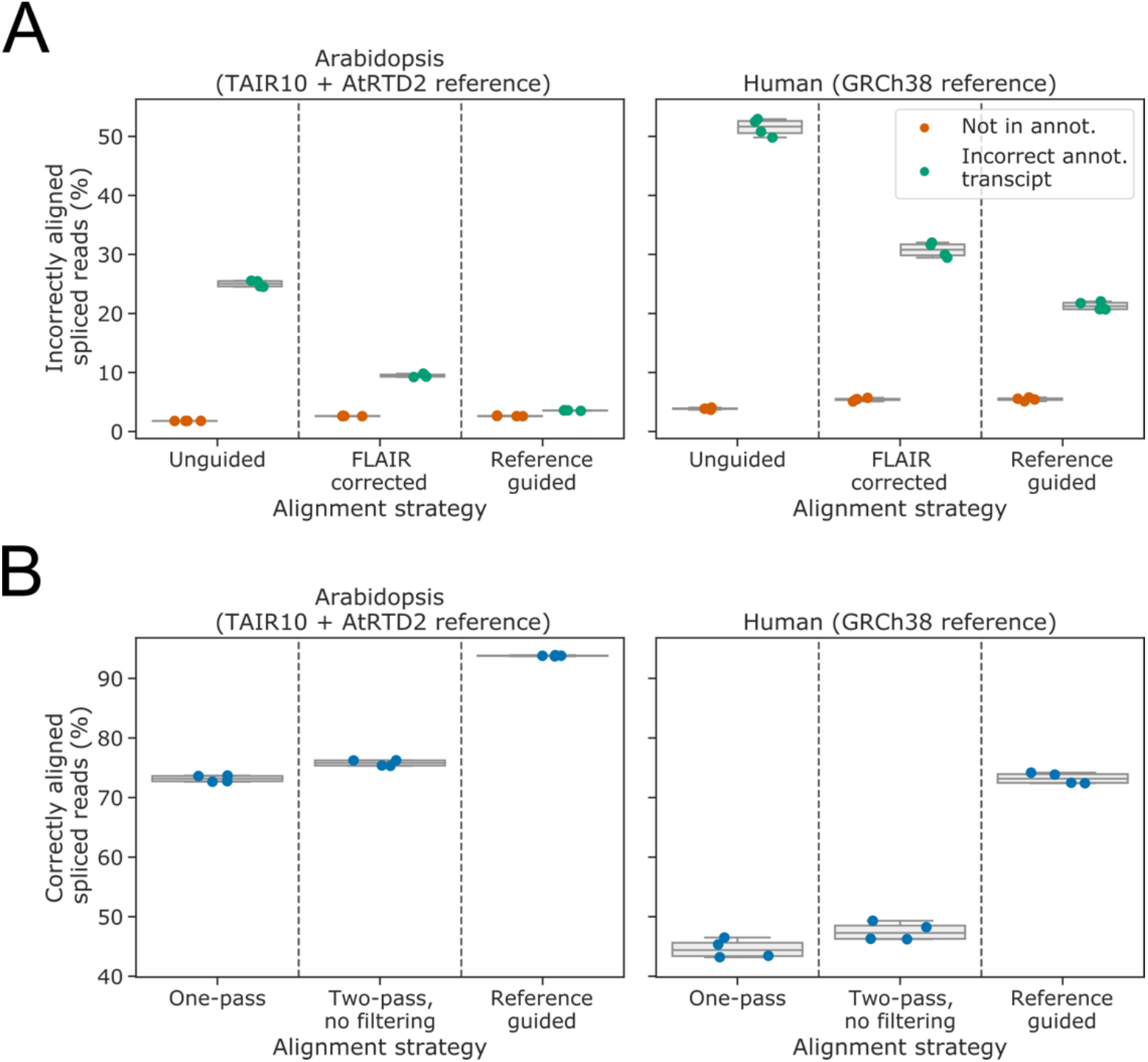
Annotation-guided alignment improves spliced alignment of simulated reads. **A** Boxplots with overlaid strip-plot showing the percentage of alignments which do not map correctly to the splice junctions of the transcript from which they were simulated, for one-pass unguided minimap2 alignments, FLAIR-corrected alignments and reference annotation-guided minimap2 alignments. Reads that align to unannotated splice junctions or combinations of junctions (“Not in annot.”) are shown in orange. Reads which align to the incorrect annotated combination of splice junctions are shown in green. Reads were simulated from Arabidopsis (left) and human (right) nanopore DRS data aligned to the AtRTD2 and GRCh38 reference transcriptomes, respectively. **B** Boxplots with overlaid strip-plot showing the percentage of alignments which map correctly to the splice junctions of the transcript from which they were simulated, for one-pass unguided minimap2 alignments, two-pass minimap2 alignment and reference annotation-guided minimap2 alignments. Reads were simulated from Arabidopsis (left) and human (right) nanopore DRS data aligned to the AtRTD2 and GRCh38 reference transcriptomes, respectively.

**Fig. S3.**
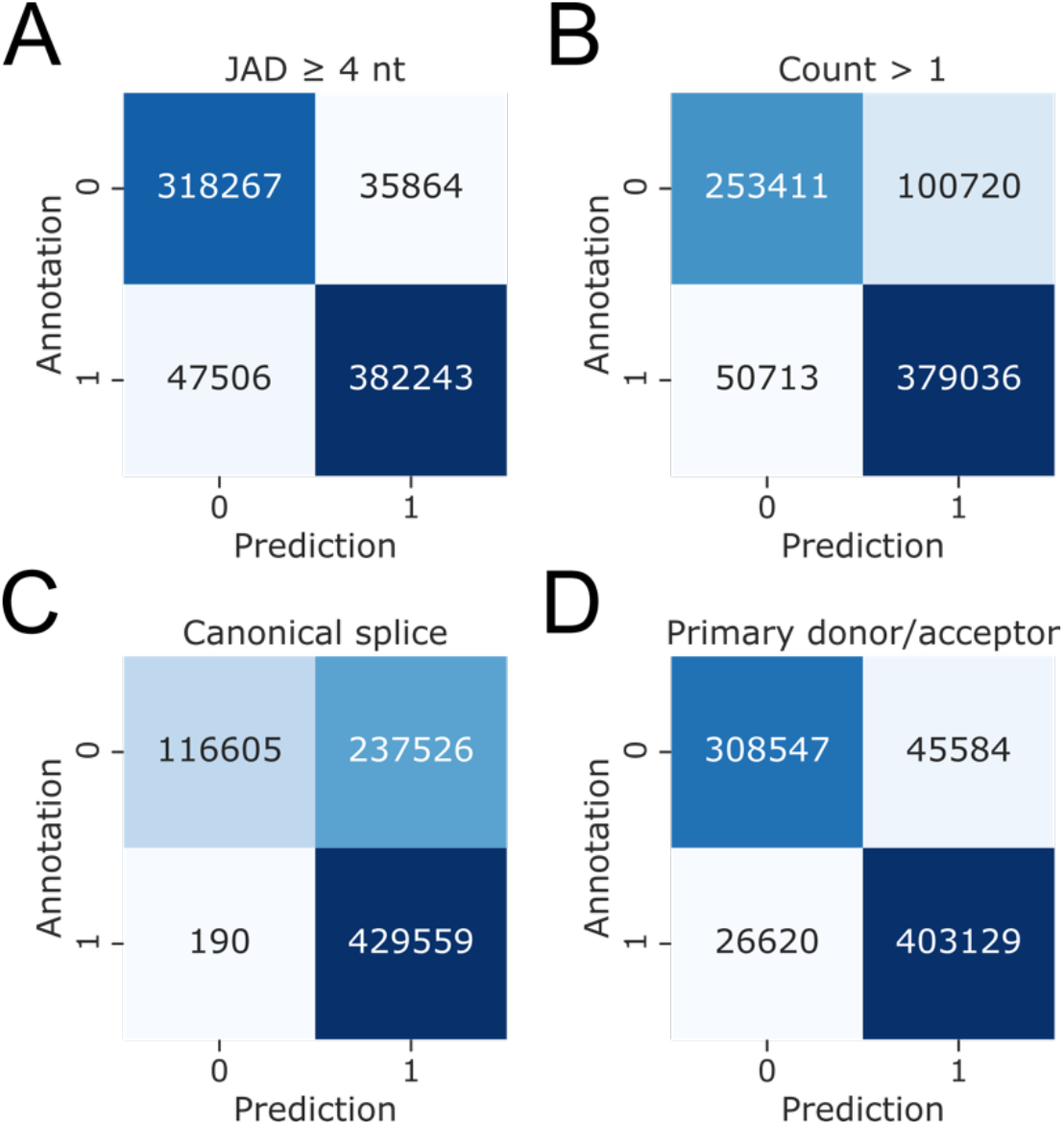
Junction metrics can identify genuine splice junctions. **A-D** Confusion matrices showing the ratios of correct and incorrect predictions using: **A** a JAD threshold of 4 nt; **B** a count threshold of 1 nt; **C** the presence of a canonical U2 GU/AG, U12 GC/AG or U12 AU/AG intron motif; and **D** the primary donor/acceptor metric, defined as whether there are no alternate donor or acceptor sites with greater support (i.e. higher count or JAD) within 20 nt.

**Fig. S4.**
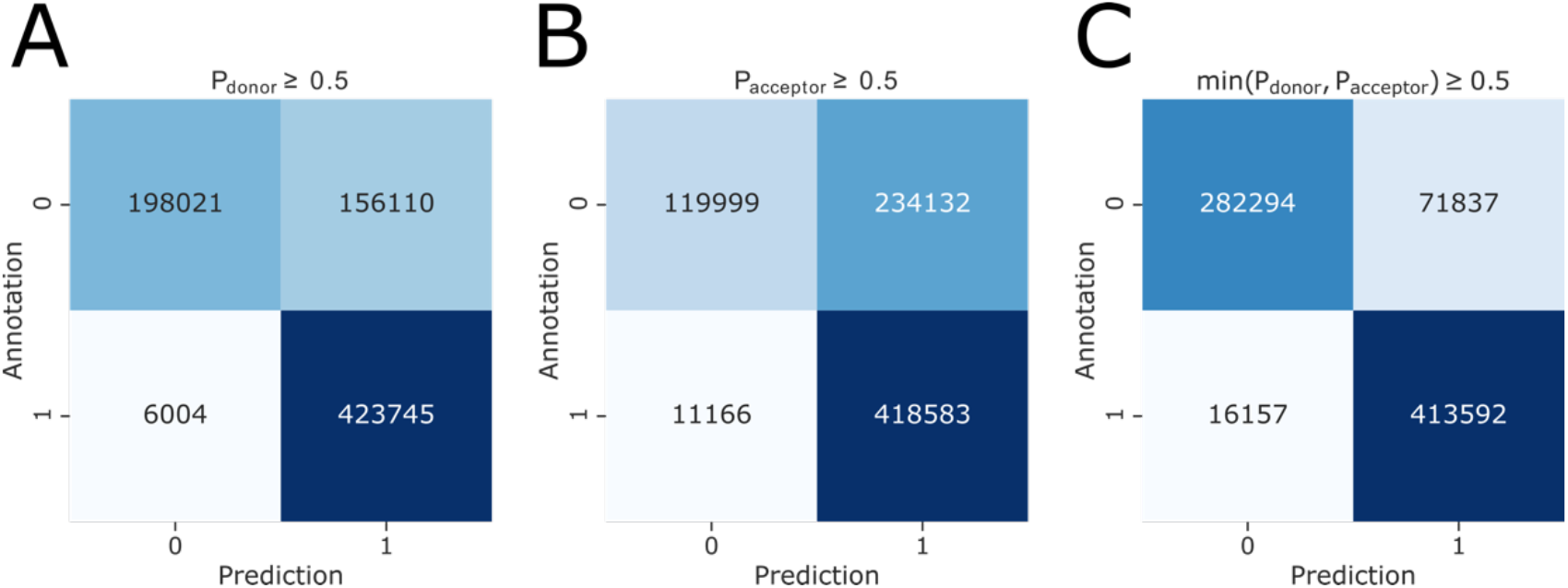
Machine-learned sequence information can identify genuine splice junctions. **A-C** Confusion matrices showing the ratios of correct and incorrect predictions using: **A** an LR prediction threshold of 0.5 for splice site strength predictions made on donor site sequences; **B** an LR prediction threshold of 0.5 for splice site strength predictions made on acceptor site sequences; **C** a minimum prediction threshold of 0.5 for both splice donor and acceptor site sequences.

**Fig. S5.**
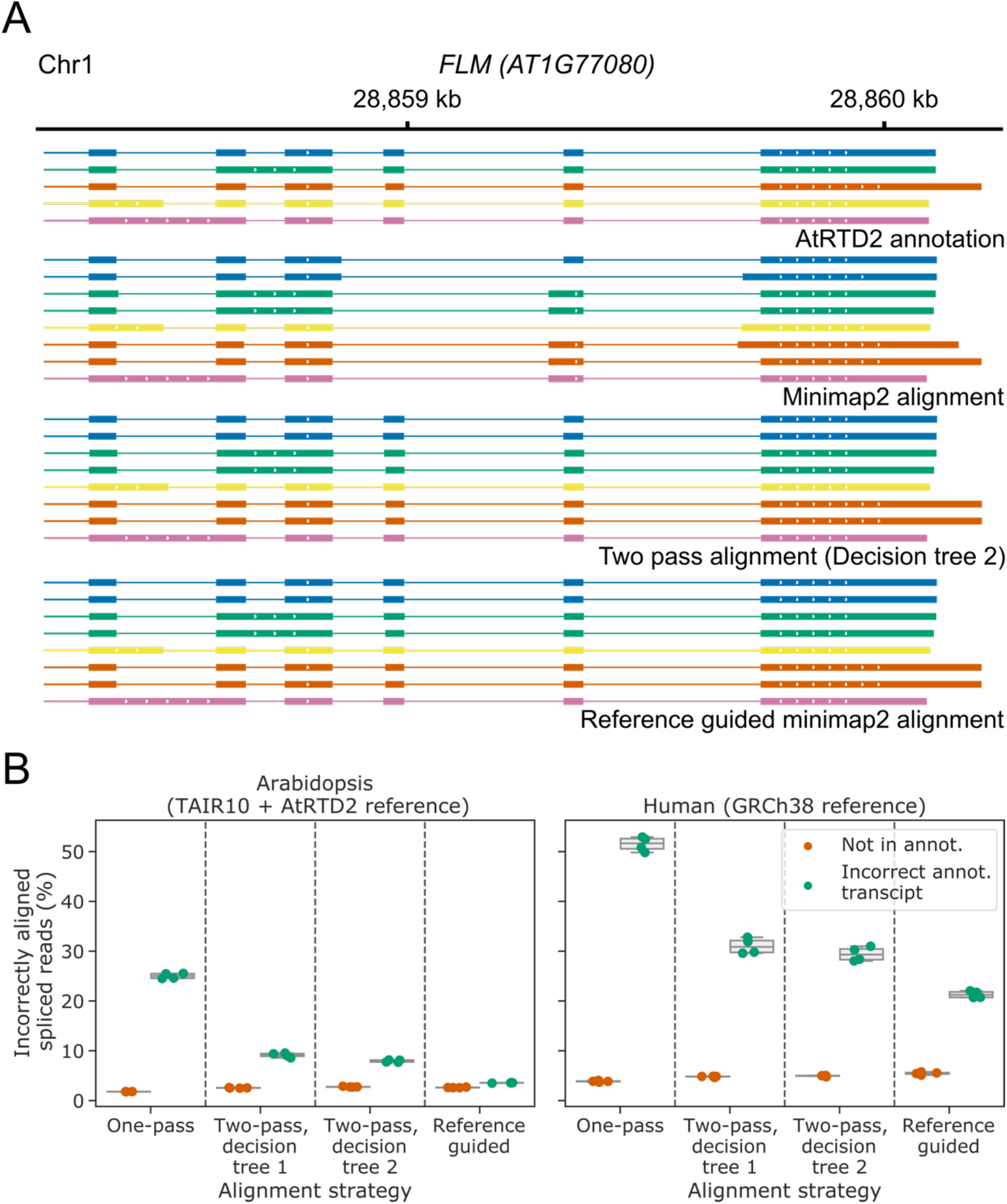
Filtered two-pass alignment improves the identification and quantification of correct transcripts without a reference annotation. **A** Gene track showing alignment of a sample of simulated nanopore DRS reads at the Arabidopsis *FLM* gene. The AtRTD2 reference annotation, from which reads were simulated, is shown on top, with unguided minimap2 alignments, two-pass minimap2 alignments using the second decision tree classification, and reference-annotation-guided alignments shown below. Only reads where exon 6 failed to align in the initial unguided alignment are shown. Each read alignment is coloured based on the reference transcript it was simulated from, and reads are in the same order within each alignment method group. Mismatches and indels are not shown. **B** Boxplots with overlaid strip-plots showing the percentage of alignments which do not map correctly to the splice junctions of the transcript from which they were simulated, for one-pass unguided minimap2 alignments, two-pass alignment with decision trees one and two, and reference annotation-guided minimap2 alignments. Reads that align to unannotated splice junctions or combinations of junctions are shown in orange. Reads which align to annotated combinations of splice junctions which they were not simulated from are shown in green. Reads were simulated from Arabidopsis (left) and human (right) nanopore DRS data aligned to the AtRTD2 and GRCh38 reference transcriptomes, respectively.

**Fig. S6.**
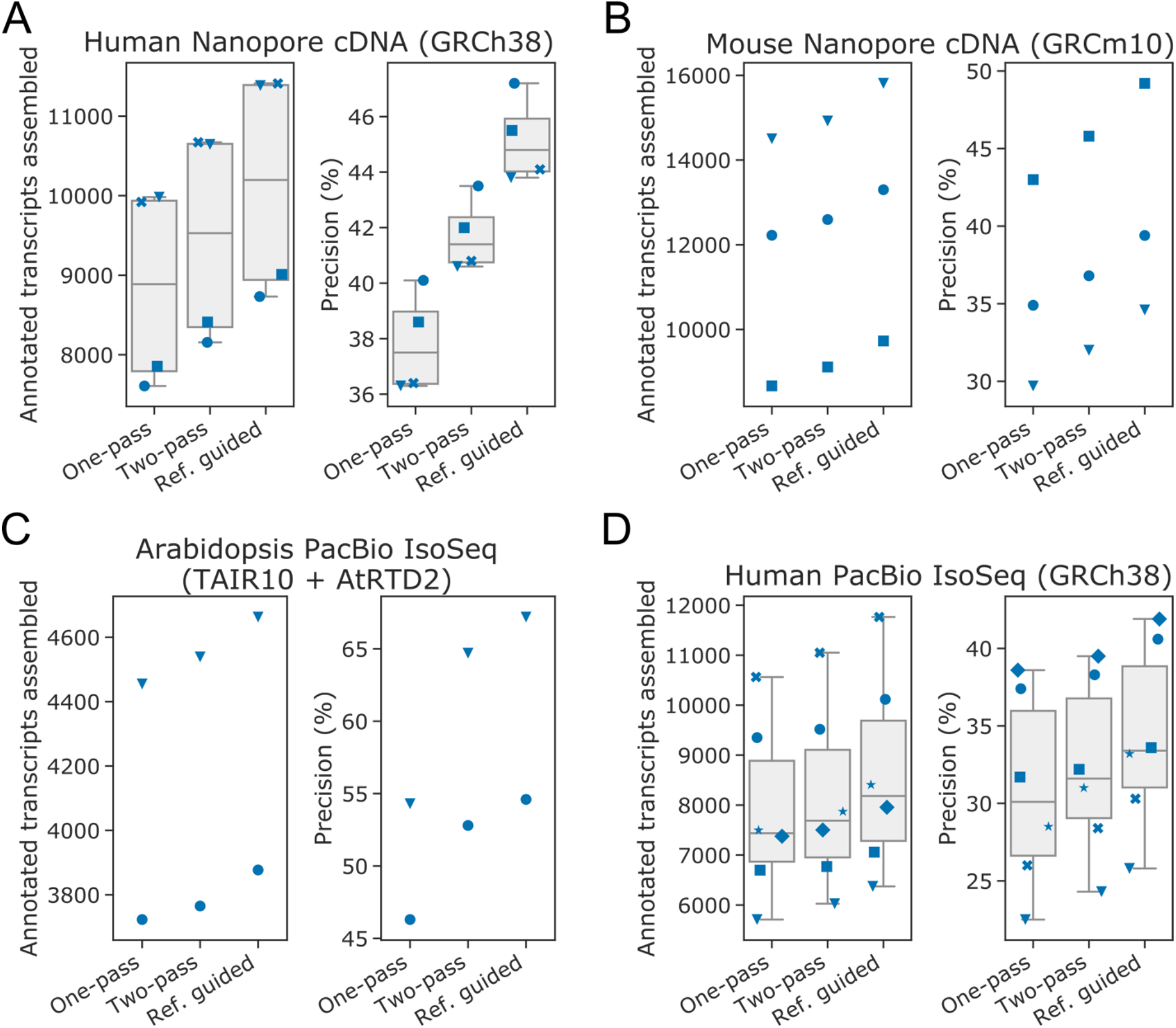
Filtered two-pass alignment improves genome-guided annotation. **A–D** Stripplots with box-and-whiskers showing the number of correct transcripts assembled (left panels) and precision of transcripts assembled (right panels) for genome-guided transcriptome assembly using StringTie2. Two-pass alignment improved the precision and number of transcripts assembled from **A** human nanopore cDNA; **B** mouse nanopore cDNA; **C** Arabidopsis PacBio IsoSeq; and **D** human PacBio IsoSeq data. For all boxplots, overlaid strip-plots are shown for individual samples. Each sample was assigned a unique marker so that changes in the metrics could be tracked between the one-pass, two-pass and reference-guided alignments. Box-and-whiskers not shown for samples with less than 4 data points.

## References

1. Wang ET, Sandberg R, Luo S, Khrebtukova I, Zhang L, Mayr C, et al. Alternative isoform regulation in human tissue transcriptomes. Nature. 2008;456(7221):470–6.

2. Stark R, Grzelak M, Hadfield J. RNA sequencing: the teenage years. Nature Reviews Genetics. 2019;20(11):631–56.

3. Mourão K, Schurch NJ, Lucoszek R, Froussios K, MacKinnon K, Duc C, et al. Detection and mitigation of spurious antisense expression with RoSA. F1000Research. 2019;8.

4. Houseley J, Tollervey D. Apparent Non-Canonical Trans-Splicing Is Generated by Reverse Transcriptase In Vitro. PLoS ONE. 2010;5(8).

5. Zhang C, Zhang B, Lin L-L, Zhao S. Evaluation and comparison of computational tools for RNA-seq isoform quantification. BMC Genomics. 2017;18(3).

6. Kovaka S, Zimin AV, Pertea GM, Razaghi R, Salzberg SL, Pertea M. Transcriptome assembly from long-read RNA-seq alignments with StringTie2. Genome Biology. 2019;20(3).

7. Pertea M, Pertea GM, Antonescu CM, Chang T-C, Mendell JT, Salzberg SL. StringTie enables improved reconstruction of a transcriptome from RNA-seq reads. Nature Biotechnology. 2015;33(3):290–5.

8. Trapnell C, Williams BA, Pertea G, Mortazavi A, Kwan G, van Baren MJ, et al. Transcript assembly and quantification by RNA-Seq reveals unannotated transcripts and isoform switching during cell differentiation. Nature Biotechnology. 2010;28(5):511–5.

9. Garalde DR, Snell EA, Jachimowicz D, Sipos B, Lloyd JH, Bruce M, et al. Highly parallel direct RNA sequencing on an array of nanopores. Nature Methods. 2018;15(3):201–6.

10. Workman RE, Tang AD, Tang PS, Jain M, Tyson JR, Razaghi R, et al. Nanopore native RNA sequencing of a human poly(A) transcriptome. Nature Methods. 2019;16(12):1297–305.

11. Parker MT, Knop K, Sherwood AV, Schurch NJ, Mackinnon K, Gould PD, et al. Nanopore direct RNA sequencing maps the complexity of Arabidopsis mRNA processing and m6A modification. eLife. 2020;9.

12. Ardui S, Ameur A, Vermeesch JR, Hestand MS. Single molecule real-time (SMRT) sequencing comes of age: applications and utilities for medical diagnostics. Nucleic Acids Research. 2018;46(5):2159–68.

13. Wick RR, Judd LM, Holt KE. Performance of neural network basecalling tools for Oxford Nanopore sequencing. Genome Biology. 2019;20(3).

14. Wick RR, Judd LM, Holt KE. Deepbinner: Demultiplexing barcoded Oxford Nanopore reads with deep convolutional neural networks. PLOS Computational Biology. 2018;14(3).

15. Dehghannasiri R, Szabo L, Salzman J, Birol I. Ambiguous splice sites distinguish circRNA and linear splicing in the human genome. Bioinformatics. 2019;35(8):1263–8.

16. Dobin A, Davis CA, Schlesinger F, Drenkow J, Zaleski C, Jha S, et al. STAR: ultrafast universal RNA-seq aligner. Bioinformatics. 2013;29(1):15–21.

17. Kim D, Paggi JM, Park C, Bennett C, Salzberg SL. Graph-based genome alignment and genotyping with HISAT2 and HISAT-genotype. Nature Biotechnology. 2019;37(8):907–15.

18. Li H, Birol I. Minimap2: pairwise alignment for nucleotide sequences. Bioinformatics. 2018;34(18):3094–100.

19. Liu B, Liu Y, Li J, Guo H, Zang T, Wang Y. deSALT: fast and accurate long transcriptomic read alignment with de Bruijn graph-based index. Genome Biology. 2019;20(3).

20. Veeneman BA, Shukla S, Dhanasekaran SM, Chinnaiyan AM, Nesvizhskii AI. Two-pass alignment improves novel splice junction quantification. Bioinformatics. 2016;32(1):43–9.

21. Gatto A, Torroja-Fungairiño C, Mazzarotto F, Cook SA, Barton PJR, Sánchez-Cabo F, et al. FineSplice, enhanced splice junction detection and quantification: a novel pipeline based on the assessment of diverse RNA-Seq alignment solutions. Nucleic Acids Research. 2014;42(8):e71–e.

22. Mapleson D, Venturini L, Kaithakottil G, Swarbreck D. Efficient and accurate detection of splice junctions from RNA-seq with Portcullis. GigaScience. 2018;7(3).

23. Zhang R, Calixto Cristiane PG, Marquez Y, Venhuizen P, Tzioutziou NA, Guo W, et al. A high quality Arabidopsis transcriptome for accurate transcript-level analysis of alternative splicing. Nucleic Acids Research. 2017;45(9):5061–73.

24. Schneider VA, Graves-Lindsay T, Howe K, Bouk N, Chen H-C, Kitts PA, et al. Evaluation of GRCh38 and de novo haploid genome assemblies demonstrates the enduring quality of the reference assembly. Genome Research. 2017;27(5):849–64.

25. Rang FJ, Kloosterman WP, de Ridder J. From squiggle to basepair: computational approaches for improving nanopore sequencing read accuracy. Genome Biology. 2018;19(3).

26. Smith TF, Waterman MS. Identification of common molecular subsequences. Journal of Molecular Biology. 1981;147(1):195–7.

27. Sheth N, Roca X, Hastings ML, Roeder T, Krainer AR, Sachidanandam R. Comprehensive splice-site analysis using comparative genomics. Nucleic Acids Research. 2006;34(14):3955–67.

28. Carrillo Oesterreich F, Herzel L, Straube K, Hujer K, Howard J, Neugebauer Karla M. Splicing of Nascent RNA Coincides with Intron Exit from RNA Polymerase II. Cell. 2016;165(2):372–81.

29. Reimer KA, Mimoso C, Adelman K, Neugebauer KM. Rapid and Efficient Co-Transcriptional Splicing Enhances Mammalian Gene Expression. bioRxiv. 2020.

30. Mercer TR, Clark MB, Andersen SB, Brunck ME, Haerty W, Crawford J, et al. Genome-wide discovery of human splicing branchpoints. Genome Research. 2015;25(2):290–303.

31. Kuo RI, Cheng Y, Smith J, Archibald AL, Burt DW. Illuminating the dark side of the human transcriptome with TAMA Iso-Seq analysis. bioRxiv. 2019.

32. Sessegolo C, Cruaud C, Da Silva C, Cologne A, Dubarry M, Derrien T, et al. Transcriptome profiling of mouse samples using nanopore sequencing of cDNA and RNA molecules. Scientific Reports. 2019;9(3).

33. Spingola M, Grate L, Haussler D, Ares M. Genome-wide bioinformatic and molecular analysis of introns in Saccharomyces cerevisiae. Rna. 1999;5(2):221–34.

34. Ares M, Grate L, Pauling MH. A handful of intron-containing genes produces the lion’s share of yeast mRNA. Rna. 1999;5(9):1138–9.

35. Chen X, Lange H, Zuber H, Sement FM, Chicher J, Kuhn L, et al. The RNA Helicases AtMTR4 and HEN2 Target Specific Subsets of Nuclear Transcripts for Degradation by the Nuclear Exosome in Arabidopsis thaliana. PLoS Genetics. 2014;10(3).

36. Lewin HA, Robinson GE, Kress WJ, Baker WJ, Coddington J, Crandall KA, et al. Earth BioGenome Project: Sequencing life for the future of life. Proceedings of the National Academy of Sciences. 2018;115(17):4325–33.

37. Koster J, Rahmann S. Snakemake--a scalable bioinformatics workflow engine. Bioinformatics. 2012;28(19):2520–2.

38. Initiative TAG. Analysis of the genome sequence of the flowering plant Arabidopsis thaliana. Nature. 2000;408(6814):796–815.

39. Quinlan AR, Hall IM. BEDTools: a flexible suite of utilities for comparing genomic features. Bioinformatics. 2010;26(6):841–2.

40. Stovner EB, Sætrom P, Hancock J. PyRanges: efficient comparison of genomic intervals in Python. Bioinformatics. 2019.

41. Pedregosa F, Varoquaux G, Gramfort A, Michel V, Thirion B, Grisel O, et al. Scikit-learn: Machine Learning in Python. J Mach Learn Res. 2011;12:2825–30.

42. Pertea M, Kim D, Pertea GM, Leek JT, Salzberg SL. Transcript-level expression analysis of RNA-seq experiments with HISAT, StringTie and Ballgown. Nature Protocols. 2016;11(9):1650–67.

